# An ancient divide in outer membrane tethering systems in Bacteria

**DOI:** 10.1101/2021.05.31.446380

**Authors:** Jerzy Witwinowski, Anna Sartori-Rupp, Najwa Taib, Nika Pende, To Nam Tham, Daniel Poppleton, Jean-Marc Ghigo, Christophe Beloin, Simonetta Gribaldo

## Abstract

Recent data support the hypothesis that Gram-positive bacteria (monoderms) arose from Gram-negatives (diderms) through loss of the outer membrane (OM). However how this happened remains unknown. Considering that tethering of the OM is essential for cell envelope stability in diderm bacteria we hypothesize that its destabilization may have been involved in OM loss. Here, we present an in-depth analysis of the four main OM tethering systems across all Bacteria. We show that their distribution strikingly follows the bacterial phylogeny with a bimodal distribution matching the deepest phylogenetic cleavage between Terrabacteria (a clade encompassing Cyanobacteria, Deinococcus/Thermus, Firmicutes, etc.) and Gracilicutes (a clade encompassing Proteobacteria, Bacteroidetes, Spirochaetes, etc.). Diderm Terrabacteria display as the main system OmpM, a porin that attaches non-covalently to modified peptidoglycan or to secondary cell wall polymers. In contrast, the lipoprotein Pal is restricted to the Gracilicutes along with a more sporadic occurrence of OmpA. While Braun’s lipoprotein Lpp is largely considered as the textbook example of OM attachment, it is actually present only in a subclade of Gammaproteobacteria. We propose an evolutionary scenario whereby the last common bacterial ancestor used a system based on OmpM, which was later replaced by one based on the lipoprotein Pal concomitantly to the emergence of the Lol machinery to address lipoproteins to the OM, with OmpA as a possible transition state. We speculate that the existence of only one main OM tethering system in the Terrabacteria would have allowed the multiple emergences of the monoderm phenotype specifically observed in this clade through OmpM perturbation. We test this hypothesis by inactivating all four *ompM* gene copies in the genetically tractable diderm Firmicute *Veillonella parvula*. The resulting mutant is severely affected in growth and displays high sensitivity to OM stress. High resolution imaging and tomogram reconstructions reveal a dramatic - yet non-lethal - phenotype, in which vast portions of the OM detach, producing large vesicles surrounding multiple monoderm-like cells sharing a common periplasm. Complementation by a single OmpM rescues the phenotype to a normal cell envelope. Together, our results highlight an ancient shift in bacterial evolution involving OM tethering systems. They suggest a possible mechanism for OM loss and a high flexibility of the cell envelope in diderm Firmicutes, making them ideal models to further refine our understanding of the mechanisms involved in bacterial OM stability, and opening the way to recapitulate the monoderm/diderm transition in the laboratory.

## Introduction

The shift between monoderms and diderms is considered one of the major transitions in the evolution of Bacteria. Growing evidence that the majority of bacterial phyla appear to be diderms and that monoderms do not constitute a monophyletic clade has led to the suggestion that the outer membrane (OM) is an ancestral character that was lost multiple times in evolution (Taib et al., 2020). However, the mechanisms involved are unknown. One of them could have been a destabilization of OM attachment not immediately leading to a lethal phenotype (Megrian et al., 2020; Taib et al., 2020; Tocheva et al., 2016).

In diderm bacteria, tethering of the OM to the cell wall is of fundamental importance to preserve cell envelope integrity. OM tethering has been mostly studied in *Escherichia coli*, where at least three different systems are identified (**Figure 1**).

**Figure 1.**
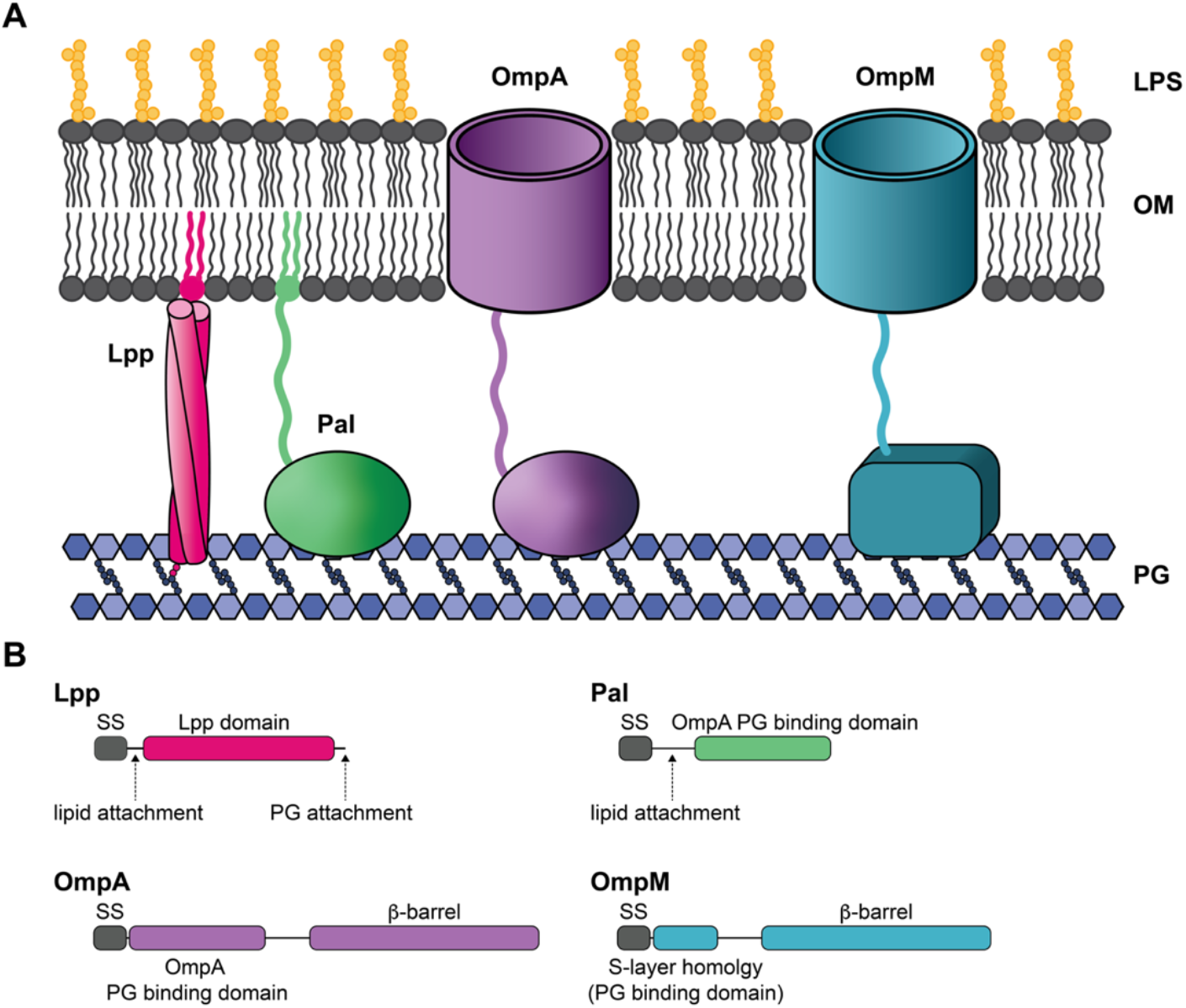
Schematic representation of the four major mechanisms for OM attachment to the cell wall (A), and domain organization of the involved proteins (B). LPS means lipopolysaccharide, OM – outer membrane, PG – peptidoglycan, SS – signal sequence.

The first to be discovered and best known is Braun’s lipoprotein (Lpp), which forms a covalent link between the OM and peptidoglycan (PG) (Braun, 1975; Braun & Rehn, 1969). Lpp mutants present increased sensitivity to membrane stress and increased OM permeability with bleb formation and hypervesiculation in minimal media (Cascales et al., 2002; Cohen et al., 2017; Suzuki et al., 1978; Yem & Wu, 1978). A second OM tethering system was discovered more recently and is based on the Pal lipoprotein, which attaches non-covalently both to PG and to the inner membrane protein TolB (Bouveret et al., 1999; Cascales et al., 2002; Parsons et al., 2006). Pal mutants display similar, yet more severe defects than Lpp mutants, including problems with OM constriction at the division plane, also considering that this protein plays multiple cellular functions (Bernadac et al., 1998; Cascales et al., 2002; Cohen et al., 2017; Gerding et al., 2007; Szczepaniak et al., 2020). In *E. coli*, the Lpp and Pal systems are at least partially redundant, as Pal overexpression is able to suppress defaults of the *lpp* null mutant (Cascales et al., 2002; Parsons et al., 2006). To our knowledge, no double mutants of Pal and Lpp have ever been produced, suggesting that at least one system has to be present for cell viability. Finally, it was demonstrated that the beta barrel protein OmpA also plays a role in OM attachment in *E.* coli (Park et al., 2012; Samsudin et al., 2017). OmpA is an integral OM protein that shares a homologous PG-binding domain with Pal, and its overexpression is able to suppress the defaults of *lpp* mutants (Cascales et al., 2002; Park et al., 2012; Samsudin et al., 2017) and the double *lpp/ompA* mutant presents a more severe phenotype than simple mutants (Bernadac et al., 1998; Sonntag et al., 1978).

Recent analyses have shown that the most ancient divide in Bacteria occurred between two large clades, the Terrabacteria (which encompass many phyla like Cyanobacteria, Firmicutes, Actinobacteria, Synergistetes, Thermus/Deinococcus, Thermotogae, Chloroflexi, the Candidate Phyla Radiation/CPR) and the Gracilicutes (including Proteobacteria, the Planctomycetes/Verrucomicrobia, Chlamydia superphylum, Nitrospirae, Nitrospina) (Coleman et al., 2021; Megrian et al., 2020, 2020). Interestingly, these data also showed that, while the Gracilicutes are very homogeneous in their cell envelope by being all diderms, the Terrabacteria include a mixture of diderms and monoderms, and that multiple losses of the OM occurred specifically in this clade (Megrian et al., 2020; Taib et al., 2020). Moreover, the Firmicutes (low GC Gram-positive bacteria) represent an ideal model to study the diderm to monoderm transition, because they comprise at least three independent clades that display an OM with lipopolysaccharide: Negativicutes, Halanaerobiales and Limnochordia (Antunes et al., 2016; Megrian et al., 2020; Taib et al., 2020; Tocheva et al., 2011). Through phylogenomic analysis, we have proposed that the OM was already present in the ancestor of all Firmicutes and would have been lost repeatedly during the diversification of this phylum to give rise to the classical monoderm cell envelope (Antunes et al., 2016; Megrian et al., 2020; Taib et al., 2020). Interestingly, diderm Firmicutes appear to lack any homologues of Lpp and Pal, but instead possess a fourth type of OM attachment (**Figure 1**). It was first characterized by *in vitro* analysis of a recombinant protein from the Negativicute *Selenomonas ruminantium* which was originally named Mep45 (Major Envelope Protein of 45 kDa) and later OmpM (Kalmokoff et al., 2009; Kamio & Takahashi, 1980; Kojima et al., 2010; Kojima & Kamio, 2012). OmpM is composed of a beta barrel that inserts into the OM and an N-terminal S-Layer Homology (SLH) domain that was shown to bind *in vitro* to PG whose peptidic chains are covalently modified with aliphatic polyamines such as cadaverine or putrescine (Kamio et al., 1981; Kamio & Nakamura, 1987; Kojima et al., 2010). OmpM homologues are present in all diderm Firmicute genomes, frequently in multiple copies and often displaying a conserved genomic context together with other proteins related to OM biogenesis and maintenance (Antunes et al., 2016; Megrian et al., 2020; Taib et al., 2020). The first proteomic characterization of the OM in the Negativicute *Veillonella parvula* DSM2008 showed that two of its four OmpM proteins are among the most abundant OM components (Poppleton et al., 2017).

Other than the Firmicutes, OmpM homologs exist in additional diderm bacterial phyla of the Terrabacterial clade. In *Synechococcus* (Cyanobacteria), it was shown that proteins composed of a periplasmic SLH domain and an OM beta-barrel (which defines OmpM) attach the OM to the PG and their depletion leads to OM stability defects (Hansel & Tadros, 1998; Kojima & Okumura, 2020). Similarly, a protein bearing the same architecture, named Slp, was identified in *Thermus thermophilus* (Deinococcus/Thermus), and its involvement in the attachment of the outer layer of the cell envelope (not yet identified at the time as a *bona fide* OM) was demonstrated (Cava et al., 2004; Fernández-Herrero et al., 1995; Olabarría et al., 1996). However, cadaverination of PG does not seem to be involved in either of these bacteria, instead the SLH domain of the OmpM homologs recognizes pyruvylated secondary cell wall polymers, and deletion of the pyruvyl transferase responsible for this modification (CsaB) led to the same phenotype as deletion/depletion of the OmpM homolog (Cava et al., 2004; Kojima & Okumura, 2020). Finally, two proteins, Ompα and Ompβ, were identified as major OM components in the Thermotogae, each corresponding to the SLH and porin domains, respectively (Engel et al., 1992), and which may represent a system derived from OmpM.

The existence of very different ways of tethering the OM in Bacteria poses the question of when they originated and how they evolved. To this aim, we explored the occurrence of these four main OM tethering systems across all Bacteria. Their distribution strikingly follows the bacterial phylogeny, with OmpM being widely present in diderm Terrabacteria, whereas Pal is restricted to the Gracilicutes along with a more scattered presence of OmpA, and the limitation of Lpp to a subclade of Gammaproteobacteria. We speculate that the existence of only one OM tethering system in the Terrabacteria would have made them more prone to OM loss, as specifically observed in this clade. To test this hypothesis, we perturbed OM tethering in the genetically tractable diderm Firmicute *Veillonella parvula* by inactivating all copies of OmpM and OmpA. The resulting mutant shows a dramatic phenotype – yet non-lethal – where vast portions of the OM detach, producing large vesicles surrounding multiple monoderm-like cells sharing a common periplasm. Complementation by a single OmpM shows a stunning capacity to rescue the phenotype and revert to a normal cell envelope. Together, our results highlight a major divide in OM tethering systems, and suggest a possible mechanism for OM loss during bacterial evolution. Diderm Firmicutes are ideal models to further our understanding of the mechanisms involved in bacterial OM stability and replay the monoderm/diderm transition in the laboratory.

## Results

### OM tethering systems display a bimodal distribution in Bacteria

We carried out an in-depth analysis of homologues of OmpM and the three other main OM attachment systems (Lpp, Pal, OmpA) across 1093 genomes representing all current bacterial phyla, including also monoderms as control. The results were then mapped onto a reference phylogeny of Bacteria, revealing that OmpM homologues are largely present in the diderm Terrabacteria (**Figure 2**, light blue dots) but are completely absent from the Gracilicutes. OmpM homologues are found in all three diderm Firmicutes clades (**Figure 2B**), suggesting they use the same mechanism to tether their OM. Moreover, Negativicutes possess a very large number of OmpM copies, up to twenty in some genomes (Supplementary Figure S1 and Supplementary Data Sheet 1C). In agreement with the experimental characterizations mentioned above, we detected OmpM homologues in the Deinococcus/Thermus, as well as Cyanobacteria and their uncultured sister clades (Melainabacteria, Blackallbacteria, Saganbacteria, and Margulisbacteria). Interestingly, Cyanobacteria have two paralogues of OmpM (Supplementary Figure S2A). One of these includes the characterized cyanobacterial OmpM and appear to be a divergent copy that might have arisen from the transfer of the SLH domain to a different porin. This copy replaced the original OmpM in multiple instances (Supplementary Figure S2A), suggesting a particular plasticity of the system in this phylum. Strikingly, we could identify OmpM homologues in all other diderm Terrabacteria phyla, including the uncultured candidate Wallbacteria, Riflebacteria, and Atribacteria, as well as in Dictyoglomi, Armatimonadetes and Synergistetes. Finally, we identified the OmpM-like system of Thermotogae (Ompα and Ompβ) in the neighbor candidate phyla (Bipolaricaulota and Fraserbacteria), suggesting that OmpM-like systems are common to the whole clade (**Figure 2**, darker blue dots).

**Figure 2.**
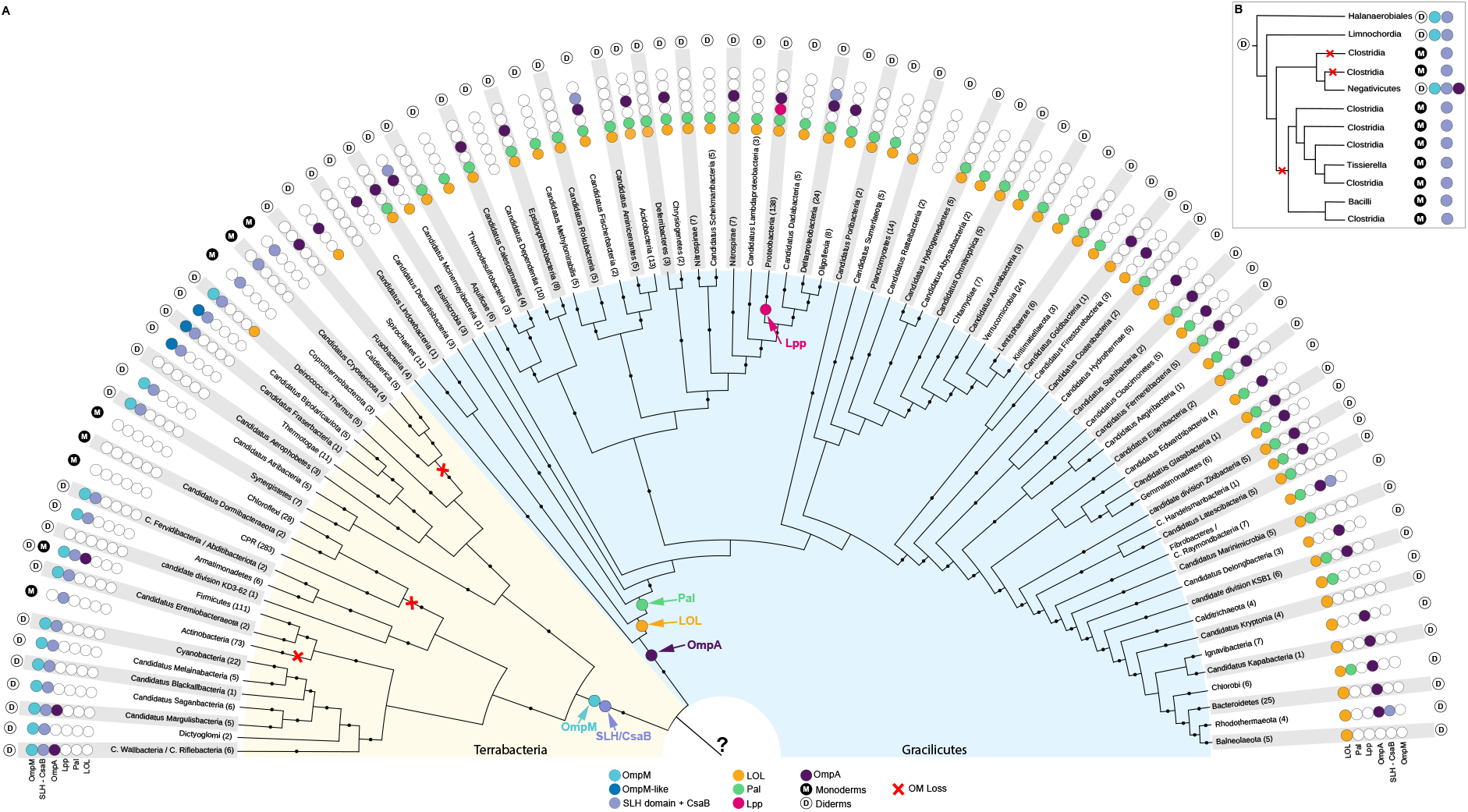
Distribution of major known OM attachment systems mapped on a reference phylogeny of Bacteria representing all current diversity (A) and on a schematic tree of Firmicutes (B). The phylogeny is based on the concatenation of RNA polymerase subunits β, β’ and elongation factor IF-2 (2,206 amino-acid positions and 377 taxa) and is rooted in between the two large clades of the Terrabacteria and Gracilicutes, according to Coleman *et al.* (2021) and Raymann *et al.* (2015). The tree was calculated using IQTREE version 1.6.3 with the model LG+C60+F+G and black dots indicate bootstrap values >90%. The presence of each of the four main attachment systems, of the Lol system and of CsaB/SLH is marked with a colored dot. Arrows indicate the possible origin of each system. The question mark in the LBCA indicates uncertainty on the presence of OmpM or OmpA (or yet another mechanism). Note that D indicates the presence of a classical OM (with or without LPS) as inferred either by experimental data or by the presence of Bam and other known systems. For this reason, Actinobacteria are marked as monoderms even if some members have an OM made of mycolic acids, which is of more recent origin. In parentheses are indicated the number of genomes analyzed. All accession numbers are given in Supplementary Data Sheet 1A and 1B.

In contrast, Pal homologues present a clearly complementary distribution, being largely present in the Gracilicutes, while they are totally absent from the Terrabacteria (**Figure 2**, green dots). The distribution of OmpA is patchier (**Figure 2**, dark violet dots), but it also concentrates mainly in the Gracilicutes. Within Terrabacteria, we could in fact find them only in some Negativicutes (including *V. parvula*), and two candidate phyla (Margulisbacteria and Wallbacteria/Riflebacteria), likely representing secondary acquisitions arising through horizontal gene transfers, as their closest homologues are nested within Gracilicutes (Supplementary Figure S2B). Finally, whereas Lpp is largely considered as the textbook example of OM tethering, our results show that it is in fact absent from the vast majority of phyla, being restricted to a subclade of Gammaproteobacteria (orders Aeromonadales, Alteromonadales, Enterobacterales, and Vibrionales) (**Figure 2** and Supplementary Data Sheet 1E), in agreement with recent publications using a much smaller taxonomic sampling (Asmar & Collet, 2018; Egan, 2018).

From these results, it appears that a major divide exists in how diderm bacteria attach their OM, with OmpM and Pal representing the main mechanisms in Terrabacteria and Gracilicutes, respectively. Because Pal is a lipoprotein, we reasoned that it may have arisen concomitantly to the emergence of the machinery to address lipoproteins to the OM, i.e. the Lol system (Braun & Hantke, 2019). Indeed, the distribution of the components of the Lol machinery on the bacterial reference tree supports our hypothesis as it shows that it is practically absent from Terrabacteria (the presence in Deinococcus/Thermus being likely due to horizontal gene transfer), while its appearance in the Gracilicutes predates that of Pal, and then closely matches its distribution (**Figure 2**, orange dots, and Supplementary Data Sheet 1A and 1B).

Finally, because the few characterized OmpM-based tethers seem to all use recognition between the SLH domain and pyruvylated secondary cell wall polymers (the use of cadaverinated PG in the Negativicutes possibly being an exception), we analyzed the concomitant distribution of homologues of CsaB, the pyruvyl transferase that is responsible for the PG modification, and of the SLH domains. Indeed, CsaB and SLH co-occur in most diderm Terrabacteria, as previously noticed by Mesnage and coworkers (Mesnage et al., 2000) on a fairly smaller sample, and match closely the distribution of OmpM, but are practically absent in the Gracilicutes (Figure 2, lilac dots). This correlation strongly suggests that OM attachment to pyruvylated PG is a conserved feature of diderm Terrabacteria. It has to be noted that we detected the presence of SLH/CsaB also in the Thermotogae and their sister phyla Bipolaricaulota and Fraserbacteria, confirming previous analyses with only one member (Mesnage et al., 2000), and suggesting that their derived OmpM-like system has maintained the same mechanism. It is interesting to add that in Bipolaricaulota CsaB is found in immediate genetic vicinity of Ompα and Ompβ.

### Inactivation of OmpM in the diderm Firmicute *Veillonella parvula* leads to growth defect and a striking vesiculated phenotype

The presence of one major OM tethering system in Terrabacteria may explain why there were multiple independent losses of the OM specifically in this clade (**Figure 2**). To test this hypothesis, we studied experimentally the consequences of perturbing OM attachment in the genetically tractable diderm Firmicute *Veillonella parvula*. The *V. parvula* SKV38 genome contains four OmpM coding genes, as well as a predicted gene coding for an OmpA homologue, but no Lpp or Pal homologues. We constructed a mutant of the three adjacent *ompM1* to *ompM3* genes (Δ*ompM1-3*), a mutant of the *ompM4* gene (Δ*ompM4*), which lies elsewhere in the genome, a quadruple deletion mutant (Δ*ompM1-4*), a mutant of *ompA* (Δ*ompA*) and a quintuple mutant (Δ*ompA*Δ*ompM1-4*). It should be noted that the Δ*ompM1-3* mutant has lost its natural competence, and we therefore successufully developed a protocol for conjugative transfert of plasmids from *E. coli* to *V. parvula*, an important addition to the genetic tools to manipulate this bacterium (see Materials and Methods). With respect to the wild type, the Δ*ompM1-3* and Δ*ompM1-4* strains presented severe growth defects, and were extremely sensitive to sodium deoxycholate, EDTA, SDS, and vancomycin (**Table 1**, Supplementary Figure S3). On the contrary, the Δ*ompM4* and Δ*ompA* mutants were not affected, suggesting they are not the main players in OM attachment in *V. parvula* (**Table 1**, Supplementary Figure S3). The quintuple Δ*ompA*Δ*ompM1-4* mutant had the strongest phenotype (**Table 1**, Supplementary Figure S3), possibly due to the fact that it lacks five of the six porins predicted in the genome (the sixth being FNLLGLLA_00833, not involved in OM attachment), which likely impairs the uptake of nutrients. Under light microscopy, the Δ*ompM1-3* cells appeared attached to large vesicles as compared to the WT (Supplementary Figure S3). The size of the vesicles was extremely variable, ranging from a few hundreds of nanometers to ~8 μm in diameter (Supplementary Figure S3). The quadruple (Δ*ompM1-4*) and quintuple (Δ*ompA*Δ*ompM1-4*) mutants presented a phenotype similar to Δ*ompM1-3*, whereas the Δ*ompM4* and Δ*ompA* mutants were not affected and their cell morphology was very similar to the WT (Supplementary Figure S3). Finally, expression of both native and HA-tagged *ompM1*, whose homologue in *V. parvula* DSM2008 corresponds to the most abundant outer membrane protein (Poppleton et al., 2017), almost completely reverted the mutant phenotype (**Table 1**, Supplementary Figure S4). However, this was not the case upon expression *in trans* of both native or HA-tagged variants of the *ompA* and *ompM4* genes (Supplementary Figure 5). Combined with the fact that there was no visible phenotype observable for the Δ*ompM4* and Δ*ompA* mutants (**Table 1**, Supplementary Figure S3), this suggests that OmpM1-3 represent the main proteins responsible for OM tethering in *V. parvula*, and that OmpA and OmpM4 have a less important role, or even a different one, such as the exchange of nutrients or/and toxic metabolic products with the environment via their porin component. We therefore focused on further analysis of the triple Δ*ompM1-3* mutant.

**Table 1.** Minimum inhibitory concentrations (MIC) of outer membrane stress inducing molecules on the five mutant strains with respect to the WT. The last four columns correspond to the complementation assay of the *ΔompM1-3* mutant. pRPF185 – empty vector. pJW34 – vector expressing native OmpM1 under the control of *tet* promoter. pJW35 – vector expressing a HA-tagged version of OmpM1 under the control of *tet* promoter. +ATc – culture realized in presence of 250 μg/l of anhydrotetracycline under chloramphenicol (25 mg/l) selection.

| Substance | WT | $\Delta ompM1-3$ | $\Delta ompM4$ | $\Delta ompA$ | $\Delta ompM1-4$ | $\Delta ompA\Delta ompM1-4$ | WT pRPF185 + ATc | $\Delta ompM1-3$ pRPF185 + ATc | $\Delta ompM1-3$ pJW34 + ATc | $\Delta ompM1-3$ pJW35 + ATc |
| --- | --- | --- | --- | --- | --- | --- | --- | --- | --- | --- |
| Sodium deoxycholate | 300 mg/l | 5 mg/l | 300 mg/l | 300 mg/l | 5 mg/l | <0.6 mg/l | 300 mg/l | 9 mg/l | 150 mg/l | 300 mg/l |
| EDTA | 2500 $\mu M$ | 156 $\mu M$ | 2500 $\mu M$ | 2500 $\mu M$ | 156 $\mu M$ | <5 $\mu M$ | 2500 $\mu M$ | 313 $\mu M$ | 1250 $\mu M$ | 2500 $\mu M$ |
| SDS | 100 mg/l | 3 mg/l | 100 mg/l | 100 mg/l | 3 mg/l | <0.2 mg/l | 50 mg/l | 3 mg/l | 25 mg/l | 50 mg/l |
| Vancomycin | >1000 mg/l | 31 mg/l | >1000 mg/l | > 1000 mg/l | 31 mg/l | <1 mg/l | 500 mg/l | 16 mg/l | 63 mg/l | 250 mg/l |

### Ultrastructural details of the triple Δ*ompM1-3* mutant

During the first manipulation of transformants, the triple Δ*ompM1-3* mutant revealed to be very delicate and easily killed by mechanical stress. 3D SIM analysis revealed further the ultrastructural defect of the Δ*ompM1-3* mutant, where the OM appeared substantially detached and formed very large vesicular structures, within which cells appeared to continue dividing (**Figure 3B** and **3C**). Although the thickness of the frozen samples (~400-500 nm) was at the limit of observable objects by Cryo-ET, we obtained an insight into the ultrastructural organization of the WT and of the Δ*ompM1-3* mutant cells preserved as close as possible to their native state, with nanometer scale resolution (**Figure 4B** and **4C**, Supplementary Figure S6), despite the fact that only a subpopulation of the triple mutant could be analyzed due to the frequent rupture of the fragile large vesicles during sample preparation. In the Δ*ompM1-3* mutant, the cells conserved their coccus shape, and were surrounded by an IM and a contiguous layer of slightly, but significatively thicker PG (15.7 ± 2.4 nm (n=30) as compared to 11.7 ± 1.8 nm (n=30) in the WT; p<10^5^ according to Student’s t test and permutation test for independent groups). Again, the OM appeared detached from the cell body, forming an enlarged, inflated periplasm with a higher density than the surrounding medium (suggesting the presence of normal proteic periplasmic components), shared by numerous dividing cells (**Figure 4B** and **4C**, Supplementary Figure S6, Supplementary Movies S2 and S3). The inner diameter of the cells was 549 ± 46 nm (n=37), not significantly different from the WT (560 ± 37 nm (n=37)). Within the vesicles, cells appeared partially attached to the OM, consistently with the 3D-SIM images. In some cases, the vesicles also contained degraded cells with patches of IM still attached to the OM (**Figure 4B**, yellow asterisk). Most of the large vesicles were connected by pearling OM tubes that could reach a few tenths of microns in length, probably remnants of budding vesicles (**Figure 4B**, white arrows). In some of the samples embedded in thinner ice, even surface pili or fimbriae still attached to the detached OM could be visualized (**Figure 4B** and Supplementary Figure S6, pink arrows). Finally, complementation of the mutant by overexpression of OmpM1 led to a normal phenotype resembling that of the WT (**Figures 3D, 4D**, Supplementary Movie S4). This indicates that a single OmpM manages to perfectly rescue such a strong phenotype, getting the OM back into place.

**Figure 3.**
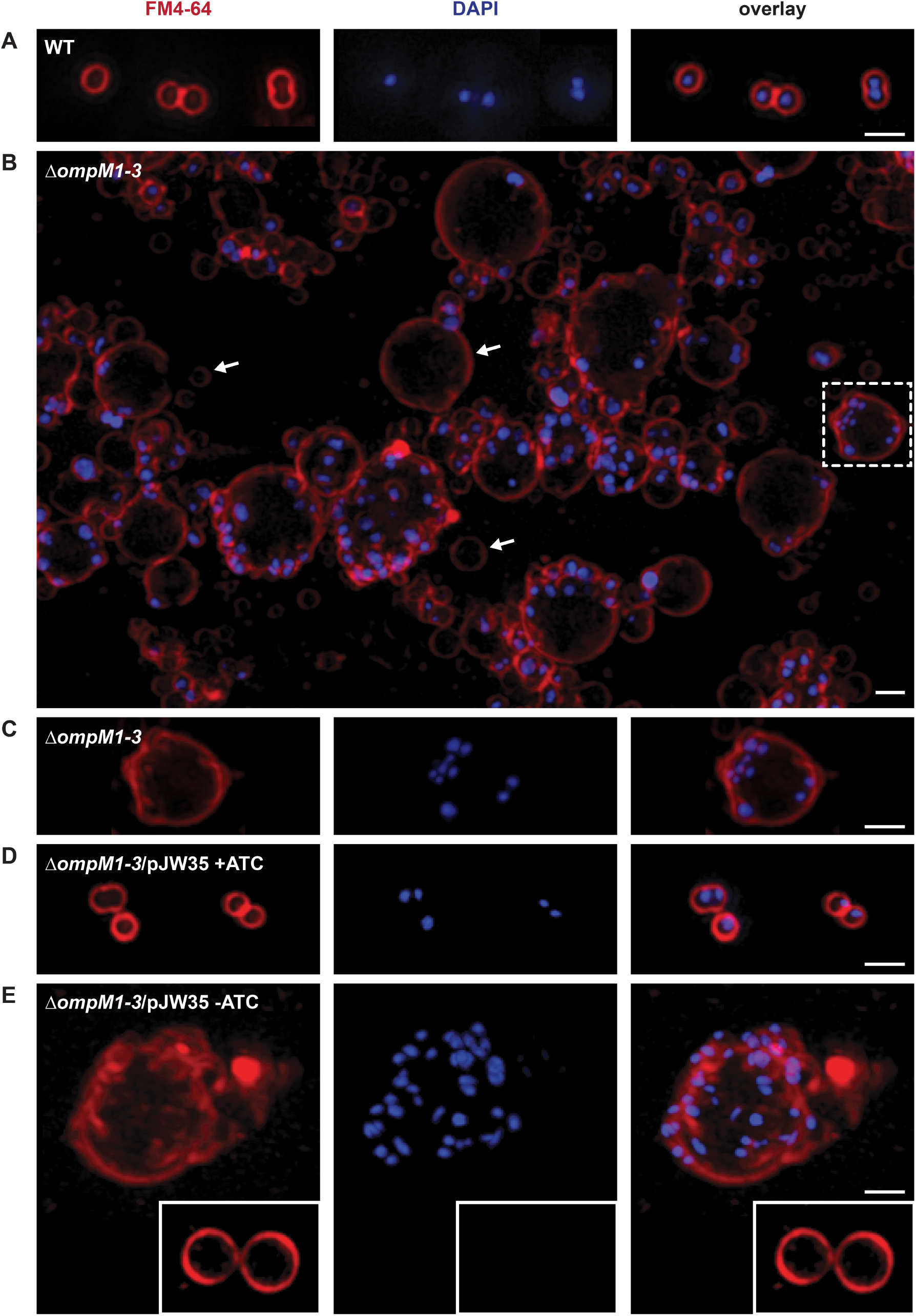
Membrane and DNA staining of representative *V. parvula* WT and Δ*ompM1-3* mutant cells imaged by 3D SIM. Cell membranes were stained with by FM4-64 (red) and the DNA by DAPI (blue). (**A**) *V. parvula* WT. (**B** and **C**) Δ*ompM1-3* mutant of *V. parvula;* the panel (**B**) presents a large field of view where arrows indicate vesicles of different sizes devoid of any DNA signal. (**D**) Δ*ompM1-3* mutant of *V. parvula* complemented with pJW35 vector expressing OmpM1-HA under control of *tet* promoter, induced overnight with 250 μg/l of anhydrotetracycline (+ATC). (**E**) Δ*ompM1-3* mutant of *V. parvula* complemented with pJW35 vector producing OmpM1-HA under control of *tet* promoter, uninduced (-ATC). Inlet shows membrane vesicles without DNA content. Scale bar in all images is 1 μm.

**Figure 4.**
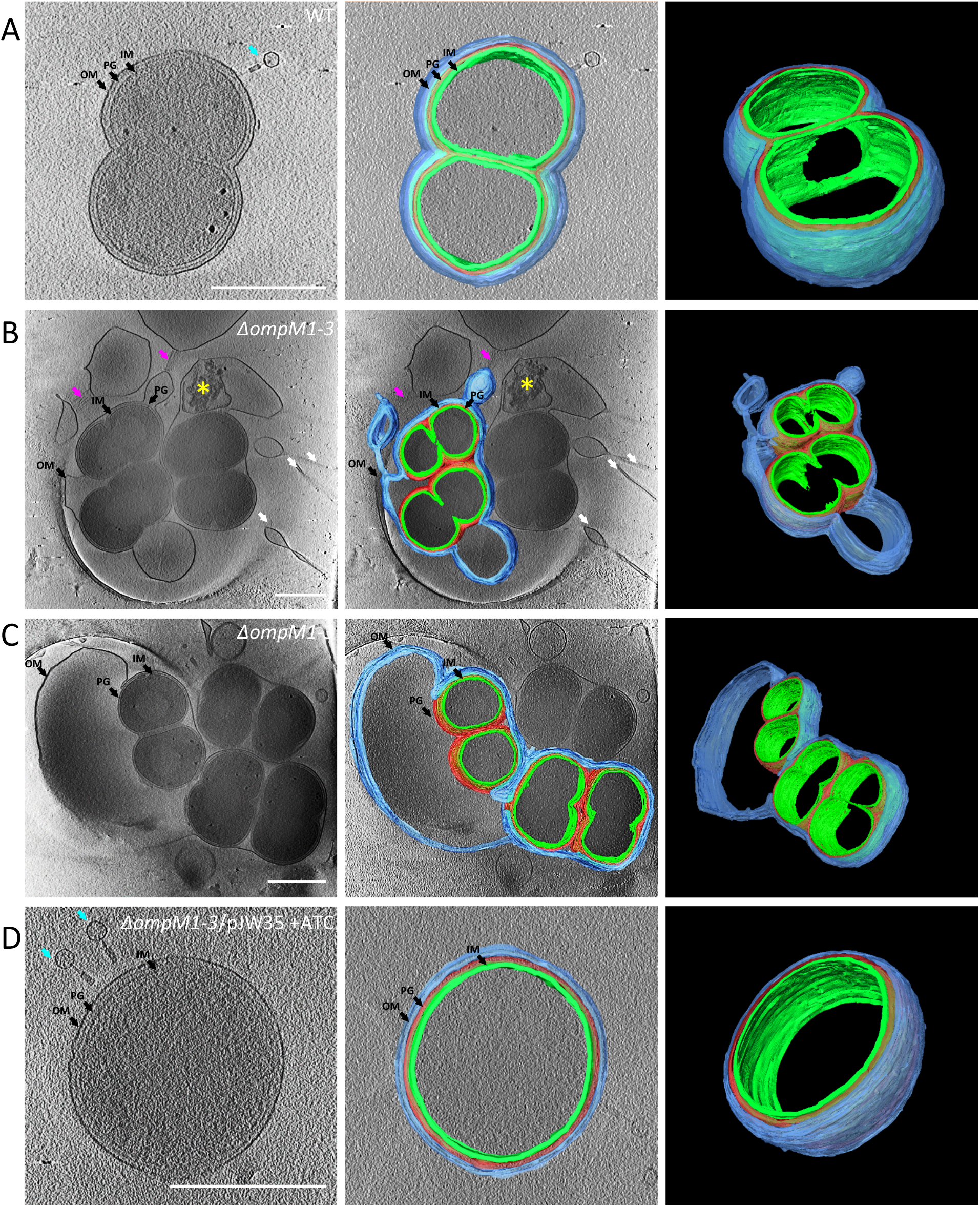
Cryo-electron tomography of *V. parvula* WT and mutant cells. (**A**) – WT, (**B-C**) – Δ*ompM1-3* mutant; detached outer membrane forming blebs and vesicles is well visible, (**D**) - Δ*ompM1*-3 mutant of *V. parvula* complemented with pJW35 vector (expressing OmpM1-HA under control of *tet* promoter) induced overnight with 250 μg/l of anhydrotetracycline. Left panels represent a sum of 10 tomogram slices (16 nm), right panels a 3D rendering of the cell envelope structure based on manual segmentation of tomogram slices, middle panels are the superposition of a tomogram slice and the 3D rendering. OM / blue marking – outer membrane, PG / red marking – peptidoglycan, IM / green marking – inner membrane. White arrows show OM tubules. Pink arrows show a bundle of fimbriae-like structures connecting two OM. Yellow asterisk shows a degraded cell. Cyan arrows show bacteriophages. All scale bars represent 0.5 μm. Videos corresponding to panels A to D, containing all slices of the 3D tomogram reconstruction are available as Supplementary Video S1 to S4.

## Discussion

In this study, we combined *in silico* and experimental approaches to reveal a major and ancient divide in the systems involved in OM stability in Bacteria, and propose a possible mechanism for the transition between diderms and monoderms.

It is possible that OmpM constituted a simple system present in the Last Bacterial Common Ancestor (LBCA) allowing OM tethering while fulfilling at the same time a generalist porin diffusion function through the membrane, as shown experimentally for the OmpM of the Negativicute *S. ruminantium* (Kojima et al., 2016) as well as for Slp of *D. radiodurans* (Farci et al., 2020). This ancestral system could then have been inherited in the branch leading to present-day Terrabacteria, and later on replaced in the branch leading to Gracilicutes by the progressive appearance of multiple redundant systems. A possible transition state would have been represented by OmpA, as it is the only system present in Fusobacteria and Spirochaetes, two phyla at the interface between Terrabacteria and Gracilicutes. The interaction of the OmpA domain to PG would have replaced that used by OmpM, making dispensable the specific PG modifications (secondary wall polymers or cadaverine) needed for SLH domain binding. Regarding the emergence of the Pal system, the origin of the Lol machinery may have represented a major event allowing the emergence of a new type of OM tethering based on lipoproteins. It has to be noted that an origin of Lol at this stage and the absence of OM lipoproteins in the LBCA is also supported by the fact that major machineries for OM biosynthesis such as Bam and Lpt in Terrabacteria do not contain OM lipoprotein components in their ancestral configuration ((Heinz et al., 2015) and personal observations). The presence of Lol and OM lipoproteins in Deinococcus/Thermus is clearly a secondary character acquired via horizontal gene transfer. Moreover, it is very likely that Pal originated from OmpA following the emergence of the Lol system, as suggested by the fact that they share the same domain to attach to PG and differ only by the mechanism of attachment to the OM. An alternative scenario would place OmpA as the OM tethering system in the LCBA, and its replacement by an OmpM-based system in the lineage leading to Terrabacteria following the appearance of pyruvylation of secondary cell wall polymers by CsaB and of SLH domains capable of recognizing it.

Whichever the ancestral OM tethering system was (OmpM, OmpA, or even some other mechanism), the reasons of its replacement by a Pal-based one are unclear. The much larger diversification of the Gracilicutes with respect to the Terrabacteria suggest that this event may have been advantageous, for instance by providing a stronger attachment allowing higher resistance to stresses on the OM. This could be the reason why the Gracilicutes did not experience any loss of the OM and are all diderms, most of them with LPS. This is in striking contrast with the Terrabacteria, whose cell envelopes are more variable, and where multiple independent losses of the OM can be inferred, at least at the divergence of three large clades of monoderms, and multiple times within the Firmicutes. It is tempting to speculate that these multiple OM losses are linked to the presence of only a single main OM attachment represented by OmpM, which could have made Terrabacteria members somehow more permissive to cell envelope perturbations.

The dramatic but non-lethal phenotype obtained by inactivating the OmpM system in *V. parvula* suggests that it may indeed have been one of the possible mechanisms for OM perturbation and eventually followed by complete OM loss. The fact that OmpM1 overexpression rescues the very severe phenotype of the triple mutant and restore a normal OM attachment indicate that this is the main involved protein, consistent with the observation that it is the most abundant of the four OmpM copies in the *V. parvula* OM proteome (Poppleton et al., 2017). The reason for the presence of a very large number of OmpM copies in the Negativicutes is unknown and remarkable, as we do not observe a similar phenomenon in other diderm Terrabacteria. This may be due to a higher tendency of these clade to recombine, also facilitated by the very close sequence similarity of these copies and the fact that they are often embedded in a conserved OM cluster, a seemingly unique feature of diderm Firmicutes (Antunes et al., 2016; Megrian et al., 2020; Taib et al., 2020).

In contrast, the lack of complementation of the *ompM1-3* deletion mutant by overexpression of OmpM4 or OmpA indicates that these proteins play a lesser role in OM attachment. Indeed, if we compare the OmpM4 SLH domain to those of OmpM1-3 a few important differences can be highlighted (Supplementary Figure S7) such as the switch of a proline from position 32 to 31 (proline being known to induce a constraint on conformation due to its secondary amine structure), the replacement of a serine (neutral amino acid) by an aspartate (positively charged amino acid) at position 41, and the replacement of an amide (asparagine or glutamine) by a hydroxyl (threonine) at position 45, might all have contributed to a loss of specificity towards cadaverinated PG.

Interestingly, we could not observe any notable difference in microscopy between the triple mutant (with only *ompM1-3* deleted) and the quintuple one (with all *ompM* and *ompA* deleted). In particular, as the majority of the cells are still attached to the inflated OM vesicles, we wondered how cells are still viable and manage somehow to keep a minimal OM attachment. Multiple envelope-spanning machineries may be involved, such as the Lpt system responsible for LPS export, or other components as for example the type IV pilus, the AcrA/AcrB/TolC transporters, or some of the recently identified autotransporter adhesins which contain an integral OM domain and an SLH domain that might anchor the protein to the PG or to secondary cell wall polymers (Béchon et al., 2020; Knapp et al., 2017; Poppleton et al., 2017).

This possible mechanism for OM destabilization likely goes beyond the Firmicutes and a few examples are available in the literature, however, in these cases, the ultrastructural details were not analyzed in depth. One concerns the *slpA* mutant in the Deinococcus/Thermus phylum, where it was observed that multiple mutant cells shared a common outermost layer of the envelope, which the authors believed to be a proteinaceous S-layer (Fernández-Herrero et al., 1995; Olabarría et al., 1996). Actually, this SlpA satisfies the criteria of an OmpM in our bioinformatic analysis and the phenotype of a *slpA* mutant is strikingly similar to that of our triple *ompM* mutant. Similar results, albeit with less severe phenotypes, involving decrease in tolerance to mechanical stress or solvents, have also been obtained for the mutant of what we know now to be an OmpM homologue, *slpA*, in *Deinococcus radiodurans*. In this case authors mention the presence of an OM as a hypothesis, instead using the term “pink envelope” (Rothfuss et al., 2006). Finally, a recent preprint confirmed that the same mechanism of OM to PG attachment is used by Cyanobacteria, as *Synechocystis* sp. PCC 6803 cells depleted for the OmpM homolog presented also a severe detachment of the OM from the cell surface (Kojima & Okumura, 2020). It is also interesting to note that both studies in *D. radiodurans* (Deinococcus/Thermus) and *S. ruminantium* (Firmicutes) noticed the occurrence of dimers and hexamers of OmpM (Farci et al., 2015; Kojima et al., 2010).

In Negativicutes, and possibly all diderm Firmicutes, binding of the SLH domain of OmpM is dependent on the presence of polyamine in the peptidoglycan peptide, cadaverine or putrescine being attached to the D-Glu residue occupying the second position in the chain (Kamio et al., 1981, 1986; Kojima et al., 2010; Kojima & Kamio, 2012). This is completely different from what is known in other Terrabacterial lineages, Cyanobacteria and Deinococcus/Thermus, where the SLH domain recognizes pyruvylated secondary cell wall polymers (Cava et al., 2004; Kojima & Okumura, 2020). Why the Negativicutes, and probably all diderm Firmicutes, changed this interaction toward cadaverinated PG remains to be understood. Future structural characterization of OmpM will certainly reveal further mechanistic insights into the functioning of this protein.

What was the fate of the OmpM system following OM loss and the multiple transitions to the monoderm phenotype in the Terrabacteria? In general, it appears that not only the porins but also the SLH domains were lost, together with the CsaB enzyme allowing the PG modification necessary for their attachment, as is clearly visible in the monoderm clade composed of the Chloroflexi, CPR, and Dormibacteraeota. In other cases, the SLH/CsaB system was kept and possibly repurposed for other functions, as is the case of a few examples within Actinobacteria, Caldiserica, Coprothermobacterota, and Ca. Cryosericota. Such repurposing is clearly visible in the Firmicutes, where the SLH/CsaB system was largely kept, despite the multiple losses of the OM. A more detailed analysis on 316 genomes covering the whole diversity of this phylum (Supplementary Data Sheet 1D) indeed shows that SLH domain and CsaB proteins co-occur (sometimes in large numbers) across all lineages. The SLH/CsaB system was mostly repurposed to specifically attach the S-layer to pyruvilated PG as described for example in *Bacillus anthracis* (Kern et al., 2010) or in *Paenibacillus alvei* (Hager et al., 2018), but also to anchor other proteins (Cann et al., 1999). The change in specificity of the SLH domain of OmpM to attach to cadaverinated PG might have perhaps allowed an easier repurposing of other SLH proteins specifically in this phylum. To confirm this hypothesis, it will be important to analyze the PG composition and the specificity of the OmpM SLH domains in the two other diderm lineages of Firmicutes – the Halanaerobiales and Limnochordia, but also investigate the specificity of the SLH domains of other proteins (for example the adhesins) present in Negativicutes.

Our results clearly demonstrate that OmpM and Pal can be considered as markers of diderm Terrabacteria or Gracilicutes, respectively. As such, their presence in newly reconstructed genomes from uncultured bacterial lineages may facilitate their taxonomic assignment. Moreover, their presence/absence may be used to infer a diderm or monoderm phenotype in the vast majority of phyla for which cell envelope characterization is missing. For example, the absence of OmpM from the Chloroflexi suggests that the diderm-like envelopes recently observed in members of this phylum may actually be other types of structures (Gaisin et al., 2020).

The lack of any predicted OM attachment system in some genomes is intriguing. While in many cases this occurs in uncultured candidate phyla and may be due to genome incompleteness, two cases are particularly interesting. For example, we could detect no known OM tethering system for Planctomycetes. Members of this phylum have a particular variant of the classic diderm cell envelope and for a long time it was believed that they lacked peptidoglycan (van Teeseling et al., 2015). Their cell membrane is also unusually dynamic, thanks to the presence of membrane coat proteins of eukaryotic type (Boedeker et al., 2017). It is possible that due to their particularities, they either evolved a novel attachment system, or so profoundly modified the one inherited from their ancestor, that it cannot be recognized anymore. Similarly, we could not detect any homologues of the four OM tethering systems in all ten available genomes from the Candidatus phylum Dependentiae. Although uncultured, members of this phylum seem to all share a similar lifestyle as intracellular symbionts of diverse amoebaes (Deeg et al., 2019; Yeoh et al., 2016), which may have led to alternative ways to tether the OM.

Our results demonstrate that Lpp, while generally considered as the textbook example of OM tethering, is in fact a very recent evolutionary invention being present in four orders within the Gammaproteobacteria (Aeromonadales, Alteromonadales, Enterobacterales, and Vibrionales; Supplementary Data Sheet 1E) and is completely absent in the rest of diderm Bacteria. This exemplifies the changes of paradigm that lay ahead with further study of the real diversity of bacterial cell envelopes. Indeed, two completely novel OM attachment systems have been recently described in Proteobacteria involving porins (Godessart et al., 2021; Sandoz et al., 2021; Wang et al., 2021): a two component system in which a periplasmic protein PapA attaches both to the peptidoglycan and to the integral outer membrane beta barrel protein OmpC, common in Betaproteobacteria (Wang et al., 2021), and numerous beta barrel outer membrane proteins covalently attached to the peptidoglycan in Alphaproteobacteria but also in some Gammaproteobacteria (Godessart et al., 2021; Sandoz et al., 2021). These results indicate that we are still far from having a complete picture of the real diversity of these mechanisms, and that further exploration is clearly needed covering the large majority of understudied bacterial phyla (including the uncultured). The combination of bioinformatics and high-resolution microscopy is a powerful approach that will surely bring unprecedented information on the real diversity of bacterial envelopes, the multiple processes involved in their biogenesis, and the mechanisms employed for maintaining OM integrity, an area of intense interest to fight bacterial pathogens.

Finally, our study indicates that *V. parvula* represents a model of choice to study the diderm/monoderm transition. First, it is one of the few genetically manipulable representatives of diderm Terrabacteria, and is a member of the Firmicutes, a phylum where multiple OM losses have occurred. Second, it has a single main OM-tether system, whose perturbation does not lead to cell death and whose strong phenotype is easily rescued by overexpression of one OmpM copy. This makes *V .parvula* ideally amenable to dissect the mechanisms involved in OM stability, with respect to more classical Gracilicutes models such as *E. coli*, which have an arsenal of -mostly essential-tethering systems and where no similar phenotypes have been observed. The OmpM triple mutant described here opens the way to further experimental work that may allow to recapitulate the diderm-to-monoderm transition in the laboratory.

### Material and methods

#### Homology searches

We assembled a databank of 1,093 genomes representing all bacterial phyla present at the National Center for Biotechnology (NCBI) as of April 2020 (for a list of taxa see Supplementary Data Sheet 1) and queried it for the presence of Lpp, Pal, OmpA, OmpM, as wells as for the Lol system and CsaB homologues.

To identify Lpp and CsaB homologues, we used HMMSEARCH from the HMMER package (Johnson et al., 2010), and screened the databank using the Pfam domains LPP (PF04728) and PS_pyruv_trans (PF04230) with the option --cut_ga. We also searched for Lpp homologues in a specific databank of 1083 proteobacteria containing 394 genomes from Gammaproteobacteria.

For the Lol system, we started by searching LolA in the databank using the Pfam domain PF03548 and HMMSEARCH with the option --cut_ga. As it is known that in some taxa LolA might be absent while the other components are present, we also searched for the ABC transporters LolC, LolD and LolE in a reduced databank composed of genomes from 192 taxa representing 36 main bacterial phyla. The alignments of families PRK10814 (LolC), PRK11629 (LolD) and PRK11146 (LolE) were downloaded from NCBI. HMM profiles were build using HMMBUILD from the HMMER package and used to query the reduced databank. Because ABC transporter subunits belong to large protein families, the results were curated manually using annotations, synteny, alignments and phylogeny. Results were pooled for the LolC/E components, as search outputs overlapped greatly, and only a clade within Gammaproteobacteria possessed these distinct two paralogs.

As OmpA and Pal homologues share the same OmpA domain (PF00691), a specific strategy was applied to distinguish them: we first searched for proteins containing the OmpA domain using HMMSEARCH and the --cut_ga option; the retrieved hits were then submitted to TMBB-PRED2 analysis (Tsirigos et al., 2016) to select those containing a beta barrel, and the positive matches were considered as OmpA proteins. As TMBB-PRED2 generates many false positives, in case of doubt the presence of a beta barrel was confirmed or infirmed with BOCTOPUS2 (Hayat et al., 2016). OmpA homologues were also identified in the same way in a local database of 230 Firmicutes. To identify Pal homologues, we used MacSyFinder (Abby et al., 2014) to investigate the immediate genetic context of OmpA-containing proteins for the presence of a TolB homologue using the Pfam domains TolB_N (PF04052), TolB_like (PF15869), WD40 (PF00400) and PD40 (PF07676).

OmpM were defined as proteins containing an SLH domain and a beta barrel porin. To find OmpM homologues, we therefore first screened our genome databank for the presence of proteins containing SLH domains (PF00395) with HMMSEARCH and the –cut_ga option. All the hits also containing a beta barrel were then identified by using TMBB-PRED2 and BOCTOPUS2. For Thermotogae, Bipolaricaulota and Fraserbacteria, the OM attachment system is similar to OmpM, however the SLH domain and the beta barrel are split into two adjacent proteins. To detect this particular configuration, we screened the proteins with an SLH domain for the presence of neighboring beta barrel structures. Results were manually curated using alignment, functional annotation, protein domains and phylogeny.

Curated OmpM homologues were then aligned using MAFFT (Katoh & Standley, 2013) with the linsi option and the alignment was trimmed by using trimal (Capella-Gutiérrez et al., 2009) with the option -gt 0.7. The tree was built using IQTREE (Kalyaanamoorthy et al., 2017; Nguyen et al., 2015) with the best-fit model LG+F+R7, and ultrafast bootstrap supports computed on 1000 replicates of the original dataset. OmpM sequences were then scanned for all Pfam domains using HMMSCAN from HMMER package.

Finally, the presence/absence of each attachment system was mapped onto a reference tree of Bacteria using custom made scripts and iTOL (Letunic & Bork, 2019).

#### Phylogenetic analysis

To build the reference bacterial phylogeny, we assembled a smaller database of 377 genomes by selecting five taxa per phylum. We selected 15 representative from the CPR and 46 from the Proteobacteria, as they are very diverse. Hidden Markov model (HMM)-based homology searches (with the option --cut_ga) were carried out with HMMERSEARCH by using the pfam profiles PF04997.12, PF04998.17, PF04563.15, PF00562.28 and PF11987.8 corresponding to RNA polymerase subunits β, β’ and translation initiation factor IF-2. Single genes were aligned using MAFFT with the linsi option, and trimmed using BMGE-1.1 (Criscuolo & Gribaldo, 2010) with the BLOSUM30 substitution matrix. The resulting trimmed alignments were concatenated into a supermatrix (377 taxa and 2,206 amino acid positions). The ML tree was generated using IQTREE v.1.6.3 with the profile mixture model LG+C60+F+G, with ultrafast bootstrap supports calculated on 1,000 replicates of the original dataset. Curated OmpA homologues were aligned using MAFFT with the linsi option and the alignment trimmed with trimal with the option -gt 0.5. The tree was built using IQTREE with the best-fit model LG+F+G4 and ultrafast bootstrap supports computed on 1000 replicates of the original dataset.

For the detailed analysis of the distribution of OmpM in the Negativicutes, we assembled a local database of 135 Negativicute proteomes, and we applied the same strategy as outlined above. Results were plotted onto a reference phylogenetic tree of Negativicutes based on a concatenation of translation initiation factor IF-2, and RNA polymerase subunits β and β’ assembled as above. Sequences were aligned using MAFFT, trimmed using BMGE, concatenated (3027 positions per sequence in final alignment) and the tree was generated using IQTREE with the best fit LG+R5 model.

For the detailed analysis of the distribution of SLH domains and CsaB proteins in Firmicutes a local database of 230 Firmicutes was queried with HMMER package using the HMM profiles for SLH (PF00395) and PS_pyruv_trans (PF04230) domains downloaded from pfam.xfam.org using the –cut_ga option.

#### Bacterial strains, culture conditions and strain manipulation

Bacterial strains used in this work are listed in Supplementary Table S1. *Escherichia coli* strains were genetically manipulated using standard laboratory procedures (Green & Sambrook, 2012). When needed, the following compounds were added to *E. coli* cultures at the following concentrations: ampicillin (liquid media) or ticarcillin (solid media) – 100 mg/l, chloramphenicol – 30 mg/l (liquid media) or 25 mg/l (solid media), apramycin – 50 mg/l, diaminopimelic acid – 300 μM, anhydrotetracycline – 250 μg/l.

*V. parvula* was manipulated as described previously (Béchon et al., 2020; Knapp et al., 2017), the culture media being either BHILC (Béchon et al., 2020) or SK (Knapp et al., 2017). When needed, the following compounds were added to *V. parvula* cultures at the following concentrations: chloramphenicol – 25 mg/l, tetracycline – 9 mg/l, erythromycin – 200 mg/l, anhydrotetracycline – 250 μg/l. The anaerobic conditions were generated using the GenBag Anaer generator (Biomérieux), the C400M anaerobic chamber (Ruskinn) or the GP Campus anaerobic chamber (Jacomex). The anaerobic chambers were filled with a H_2_ /CO_2_/N_2_ (5%/5%/90%) mixture.

#### Plasmids, primers and DNA manipulations

All plasmids and primers used in this study are listed in Supplementary Table S2 and S3, respectively. Clonings were performed using NEBuilder HiFi DNA Assembly Master Mix (New England Biolabs). Chemocompetent homemade *E. coli* DH5α cells (Inoue et al., 1990) were used for transformation of cloning products or plasmids. *V. parvula* genomic DNA was extracted according to a protocol previously described for *Streptomyces* gDNA extraction (Jacques et al., 2015) from stationary phase cultures in SK medium.

#### Conjugation between *E. coli* and *V. parvula*

The plasmid to conjugate was introduced into *E. coli* MFDpir strain by electrotransformation using standard protocol (Robey et al., 1996). An overnight stationary culture was used to start a fresh culture by 1/100 dilution (180 rpm, 37°C, LB medium). When the culture reached OD_600_ comprised between 0.4 and 0.9, the cells were washed twice by an equal volume of LB, resuspended in 1/10 of initial volume and stored on ice upon further use. If for any reason the conjugation could not be carried out the same day, the prepared *E. coli* was stored in 4°C for up to a week. *V. parvula* cultures were launched from cells scratched from a SK petri dish in SK liquid medium at initial OD_600_ of 0.03 and grown anaerobically at 37°C until the OD reached 0.1 – 0.2. 0.5 ml of donor *E. coli* strain was mixed with 1 ml of acceptor *V. parvula* strain, centrifugated briefly, the excess volume of medium removed, and the resuspended cells were deposited in form of a droplet on an SK medium plate containing diaminopimelic acid. After 24h of anaerobic incubation at 37°C, the cells were scratched from the Petri dish, resuspended in SK medium and plated on selective SK medium without diaminopimelate.

#### Generation of *ompM* and *ompA* deletion mutants

The mutants were generated using the technique described previously (Knapp et al., 2017). Briefly, for *ompM4* and *ompA* deletion upstream and downstream fragments of the genes to delete as well as the chloramphenicol resistance cassette *catP* were amplified and assembled into one linear construct by PCR. For *ompM1-3* deletion, the tetracycline resistance cassette *tetM* was used, and the PCR assembly step has proven difficult, so the three fragments were cloned into pUC18 *Xba*I site, and then the linear construct was obtained by PCR amplification. Natural competence was used to transform the SKV38 WT strain with the linear construct and induce homologous recombination (and subsequent gene replacement with the resistance marker) and the correct mutant construction was verified by PCR amplification (and sequencing of amplicons) of the recombination junction zones. The details of the constructions are given in Supplementary Information.

#### Generation of quadruple and quintuple mutants

To generate a quadruple Δ*ompM1-4* mutant, the Δ*ompM4* mutant was transformed with Δ*ompM1-3* gDNA according to previously established protocol (Knapp et al., 2017). The correctness of the mutant was verified by the PCR of the recombination junction zones, and by the PCR verification of the absence of *ompM4* gene.

To generate a quintuple mutant, a novel linear construct was generated by PCR assembly, using upstream and downstream fragment of *ompA* gene, and the *ermE* resistance marker. This construct was used for transformation of *ompM4* mutant according to previously established protocol (Knapp et al., 2017), yielding Δ*ompA*Δ*ompM4* strain, which was in turn transformed by the gDNA of the Δ*ompM1-3* strain, yielding the desired Δ*ompA*Δ*ompM1-4* strain. The correct mutant construction was verified by PCR amplification (and sequencing of amplicons) of the recombination junction zones as well as the absence of *ompA* and *ompM4* genes by PCR. The details of the constructions are given in Supplementary Information.

#### OmpM1, OmpM4 and OmpA Expression vector construction

The expression vectors were constructed by cloning the gene of interest with its native RBS or with a modified RBS into *Sac*I site of pRPF185 *Escherichia/Clostridium* conjugative shuttle tetracycline inducible expression vector (Fagan & Fairweather, 2011). In some vectors, an HA-tag was added during the cloning process into a predicted extracellular loop of the beta barrel (the exact point of insertion of the HA-tag is presented for OmpM1 in the Supplementary Figure S8). The integrity of constructs was verified by sequencing. All vectors are described in the Supplementary Table S2 and all the details of the construction process are described in Supplementary Information.

#### Minimum inhibitory concentration (MIC) determination and growth curves

For MIC, serial twofold dilutions of sodium deoxycholate, EDTA, SDS and vancomycin were made in BHILC medium. The dilutions were put in 96-well flat bottom plates (150 μl per well). Bacterial strains to be tested were grown on SK plates (37°C, anaerobic conditions) for 2-3 days. They were scratched from the plate, resuspended in BHILC medium, and the OD_600_ was adjusted to 1.5. The wells were inoculated with 5 μl of bacterial suspension in order to obtain an initial OD_600_ of 0.05 and then incubated anaerobically in 37°C, 180 rpm for 24h. The MIC was defined as the lowest concentration for which the final OD was inferior to 0.1.

For growth curves, a 96-well flat bottom plate was filled with 250 μl per well of BHILC medium. The medium was partially degassed in −0.6 bar vacuum and preincubated 4h inside the GP Campus anaerobic chamber (Jacomex). Then, the wells were anaerobically seeded with 5 μl of the same bacterial suspension as for the MIC, the concentration in the well being 0.03. The plates were then sealed with Adhesive PCR Plate Seals (Thermo Fisher Scientific). They were then incubated in TECAN Sunrise spectrophotometer for 24 hours at 37 °C. OD_600_ was measured every 30 minutes, after 15 min of orbital shaking. Each test was performed in pentaplicate.

#### Cultures for microscopy observation

For *V. parvula* a liquid culture of 5 ml (in 50 ml Falcon tube) was launched with a glycerol stock in SK medium. The culture was incubated anaerobically at 37°C. For strains bearing no plasmid, the observations were made after two days of culture. As for expression vector bearing strains, at the end of the second culture day the original culture was diluted 10x in fresh medium and incubated anaerobically overnight at 37°C either in presence or in absence of anhydrotetracycline.

For *E. coli* a liquid culture of 10 ml (in 50 ml Falcon tube) was launched from a glycerol stock in LB medium. The culture was incubated aerobically 24h at 37°C with 180 rpm rotative shaking. A new culture was launched by diluting the original culture 10x in fresh anhydrotetracycline containing medium. The culture was then incubated overnight in a 250 ml Erlenmeyer flask at 20°C with 180 rpm rotative shaking.

#### 3D Structured Illumination Microscopy (3D SIM)

An aliquot of culture was mixed with FM 4-64 membrane staining dye (Thermo Fischer Scientific) at a final concentration of 5 mg/l and DAPI at a final concentration of 17 mg/l. Bacterial cell suspensions were applied on high precision coverslips (No. 1.5H, Sigma-Aldrich) coated with a solution of 0.01 % (w/v) of poly-L-lysine. After letting the cells attach onto the surface of the coverslip for 10 min, residual liquid was removed, 8 μL of antifade mounting medium (Vectashield) were applied and the coverslip was sealed to a slide. SIM was performed on a Zeiss LSM 780 Elyra PS1 microscope (Carl Zeiss, Germany) using C Plan-Apochromat 63× / 1.4 oil objective with a 1.518 refractive index oil (Carl Zeiss, Germany). The samples were excited with laser at 405 nm for the DAPI staining and 561 nm for the FM4-64 staining and the emission was detected through emission filter BP 420-480 + LP 750 and BP 570-650 + LP 750, respectively. The fluorescence signal is detected on an EMCCD Andor Ixon 887 1 K camera. Raw images are composed of fifteen images per plane per channel (five phases, three angles), and acquired with a Z-distance of 0.10 μm. Acquisition parameters were adapted from one image to one other to optimize the signal to noise ratio. SIM images were processed with ZEN software (Carl Zeiss, Germany) and then corrected for chromatic aberration using 100 nm TetraSpeck microspheres (ThermoFisher Scientific) embedded in the same mounting media as the sample. For further image analysis of SIM image z stacks we used Fiji (ImageJ) Version 2.0.0-rc-68/1.52i (Schindelin et al., 2012). Namely, we assigned a color to the fluorescent channels, stacks were fused to a single image (z projection, maximum intensity) and brightness and contrast were slightly adapted. Regions of interest were cut out and, for uniformity, placed on a black squared background. Figures were compiled using Adobe Illustrator 2020 (Adobe Systems Inc. USA).

#### Cryogenic electron microscopy observation (CryoEM)

A solution of BSA-gold tracer (Aurion) containing 10-nm-diameter colloidal gold particles was added to a fresh culture of *V. parvula* with a final ratio of 3:1. A small amount of the sample was applied to the front (4 μl) and to the back (1.2 μl) of carbon-coated copper grids (Cu 200 mesh Quantifoil R2/2, Quantifoil, or Lacey, EMS), previously glow discharged 2 mA and 1.5-1.8×10-1 mbar for 1 min in an ELMO (Corduan) glow discharge system. The sample was then vitrified in a Leica EMGP system. Briefly, the excess liquid was removed by blotting with filter paper the back side of the grids for 6-7 s at 18°C and 95% humidity, and then the sample was rapidly frozen by plunging it in liquid ethane. The grids were stored in liquid nitrogen until image acquisition in the TEM.

Cryo-electron microscopy was performed on a Tecnai 20 equipped with a field emission gun and operated at 200 kV (Thermo Fisher) using a Gatan 626 side entry cryoholder. Images were recorded using the SerialEM software on a Falcon II (FEI, Thermo Fisher) direct electron detector, with a 14 μm pixel size. Digital images were acquired at nominal magnification of 29000x, corresponding to pixel size of 0.349 nm. For high-magnification images, the defocus was −8 μm. Cell width and peptidoglycan width were measured using Fiji (Schindelin et al., 2012).

Dose-symmetric tilt series were collected on a 300kV Titan Krios (Thermo Fisher Scientific) transmission electron microscope equipped with a Quantum LS imaging filter (Gatan, with slit width of 20 eV), single-tilt axis holder and K3 direct electron detector (Gatan). Tilt series with an angular increment of 2° and an angular range of ±60° were acquired with the Tomography software (Thermo Fisher Scientific). The total electron dose was 130 electrons per Å2 at a pixel size of 8 Å. Dose symmetric tilt series were saved as separate stacks of frames and subsequently motion-corrected and re-stacked from −60° to +60° using IMOD’s function align frames (Mastronarde & Held, 2017) with the help of a homemade bash script. Three-dimensional reconstructions were calculated in IMOD by weighted back projection and a gaussian filter was used to enhance contrast and facilitate subsequent segmentation analysis. The 3D segmentation of the tomograms was performed by using the Amira software (Thermo Fisher Scientific).

## Supporting information

Supplementary Data Sheet 1

## ACKNOWLEDGEMENTS

This work was supported by funding from the French National Research Agency (ANR) (Fir-OM ANR-16-CE12-0010), the Institut Pasteur “Programmes Transversaux de Recherche” (PTR 39-16), the French government’s Investissement d’Avenir Program, Laboratoire d’Excellence “Integrative Biology of Emerging Infectious Diseases” (grant n°ANR-10-LABX-62-IBEID) and the Fondation pour la Recherche Médicale (grant DEQ20180339185). N.P. is funded by a Pasteur-Roux Postdoctoral Fellowship from the Institut Pasteur. We thank Alicia Jiménez-Fernández for help with *Veillonella* genetics techniques. We gratefully acknowledge the UTechS Photonic BioImaging (Imagopole), C2RT, Institut Pasteur (Paris, France) and the France–BioImaging infrastructure network supported by the French National Research Agency (ANR-10–INSB–04; Investments for the Future), and the Région Ile-de-France (program Domaine d’Intérêt Majeur-Malinf) for the use of the Zeiss LSM 780 Elyra PS1 microscope. We thank S. Tachon from the NanoImaging Core facility of the Center for Technological Resources and Research of Institut Pasteur for asistance with the tomography acquisitions at the Titan Krios microscope. We are grateful for equipment support from the French Government Programme Investissements d’Avenir France BioImaging (FBI, N° ANR-10-INSB-04-01). We thank M. Nilges and the Equipex CACSICE (Centre d’analyse de systèmes complexes dans les environnements complexes) for providing the Falcon II direct detector. The authors acknowledge the IT department at Institut Pasteur, Paris, for providing computational and storage services (TARS cluster).

## AUTHORS CONTRIBUTIONS

JW performed all molecular biology and microbiology experiments with the assistance of TNT. ASR and JW performed the electron microscopy data acquisition, reconstructions and visualizations, and NP performed the 3D-SIM data acquisition and treatment. NT and JW performed the bioinformatic and evolutionary analysis. DP performed preliminary bioinformatic analyses in an early version of the manuscript. JMG provided lab facilities. CB and SG supervised the study. JW, CB and SG wrote the paper with contributions from ASR, NT, NP, and JMG. All authors contributed to the final version of the manuscript.

## Notes

https://drive.google.com/drive/folders/1xpQGI-u-5WeNOQrs5mCSIj9urhfaJjfn?usp=sharing

### Competing Interest Statement

The authors have declared no competing interest.

