## Supplementary Information for "An ancient divide in outer membrane tethering systems in Bacteria"

### An ancient divide in outer membrane tethering systems in Bacteria – SUPPLEMENTARY INFORMATION

#### Supplementary Figures

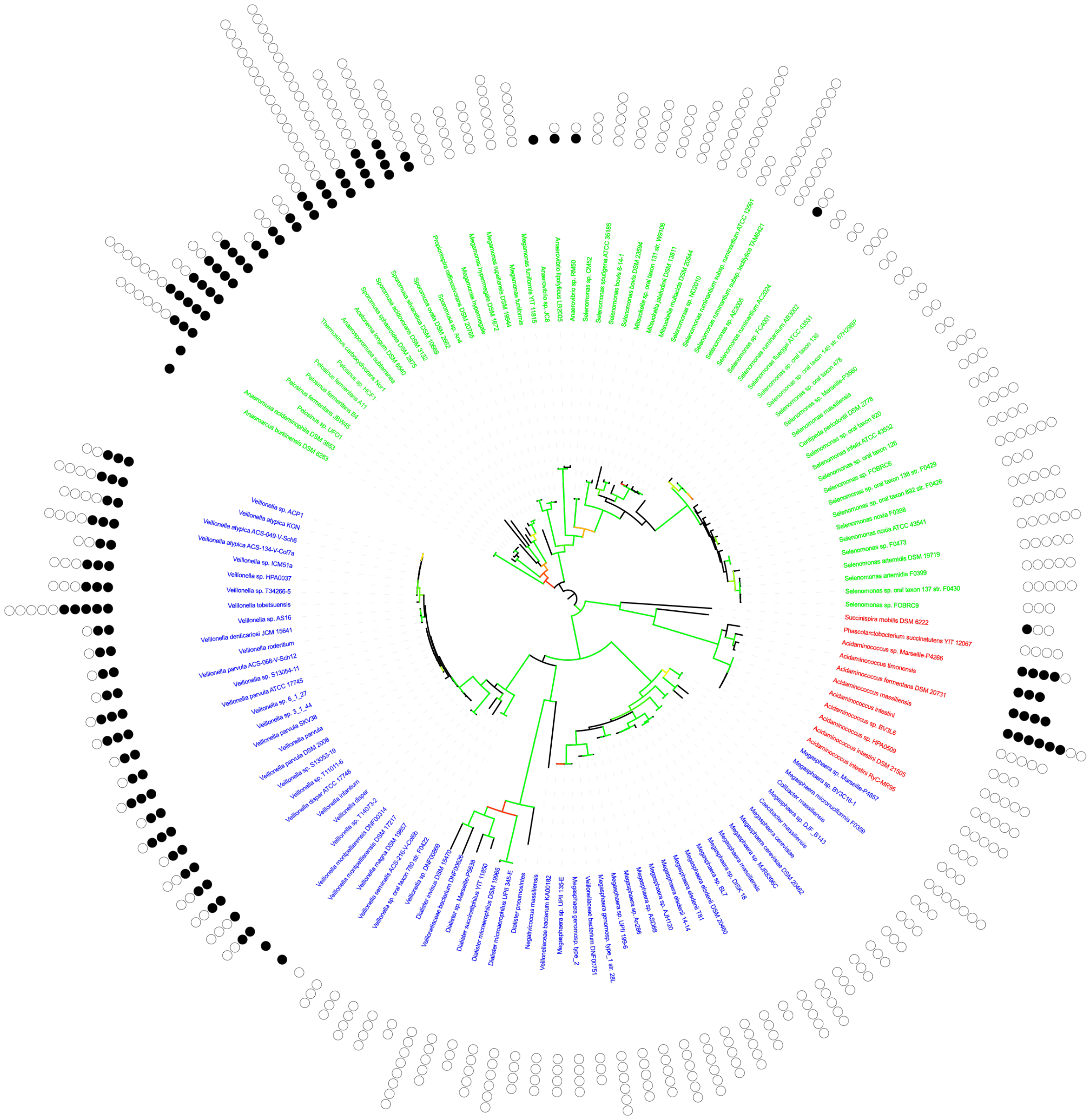

**Supplementary Figure S1. Number of *ompM* copies per genome plotted on a reference Negativicutes tree.** Selenomonadales are labelled in green, Acidaminococcales in red and Veillonellales in blue. The phylogeny is based on the concatenation of RNA polymerase subunits β, β′ and elongation factor IF-2 (3,027 amino-acid positions) The tree was calculated using IQTREE version 1.6.3 with the model LG+R5. Colours of branches represent bootstrap values from 80 (red) to 100 (green). The *ompM* copies present inside the OM gene cluster are marked by full circles, those outside the cluster by empty ones.

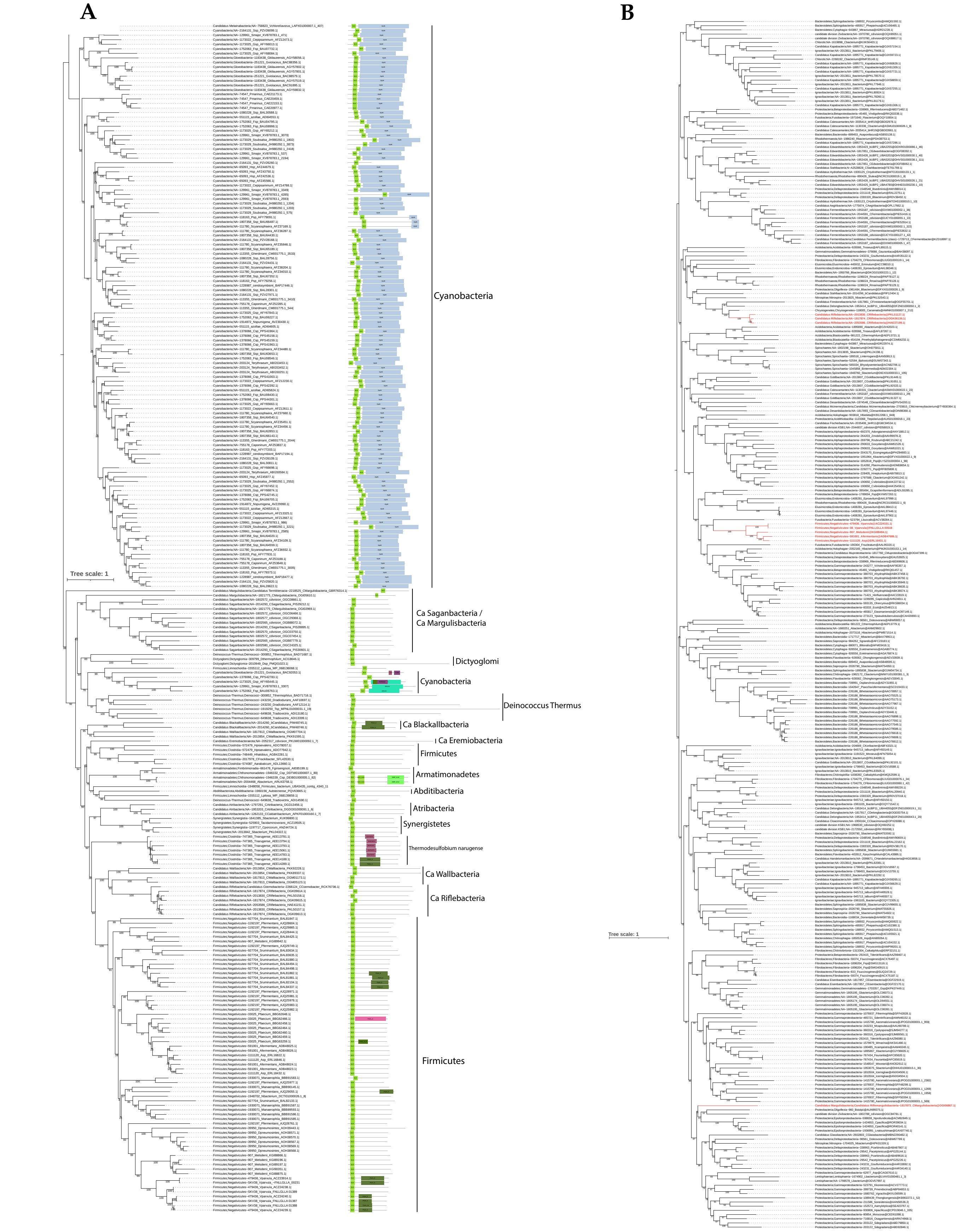


**Supplementary Figure S2. (A) Phylogeny of OmpM homologues in bacteria with the organization of identified pfam domains. (B) Phylogeny of OmpA homologues in bacteria.** Maximum likelihood tree built from an alignment of 261 (OmpM) / 289 (OmpA) sequences and 367 aa (OmpM) / 371 (OmpA) positions. Supports at the nodes correspond to ultrafast bootstraps. The scale bar corresponds to the average number of substitutions per site. The identified pfam domains were plotted using IToL (Letunic and Bork, 2019) on (A), and on (B) the sequences retrieved from Terrabacteria are marked in red.

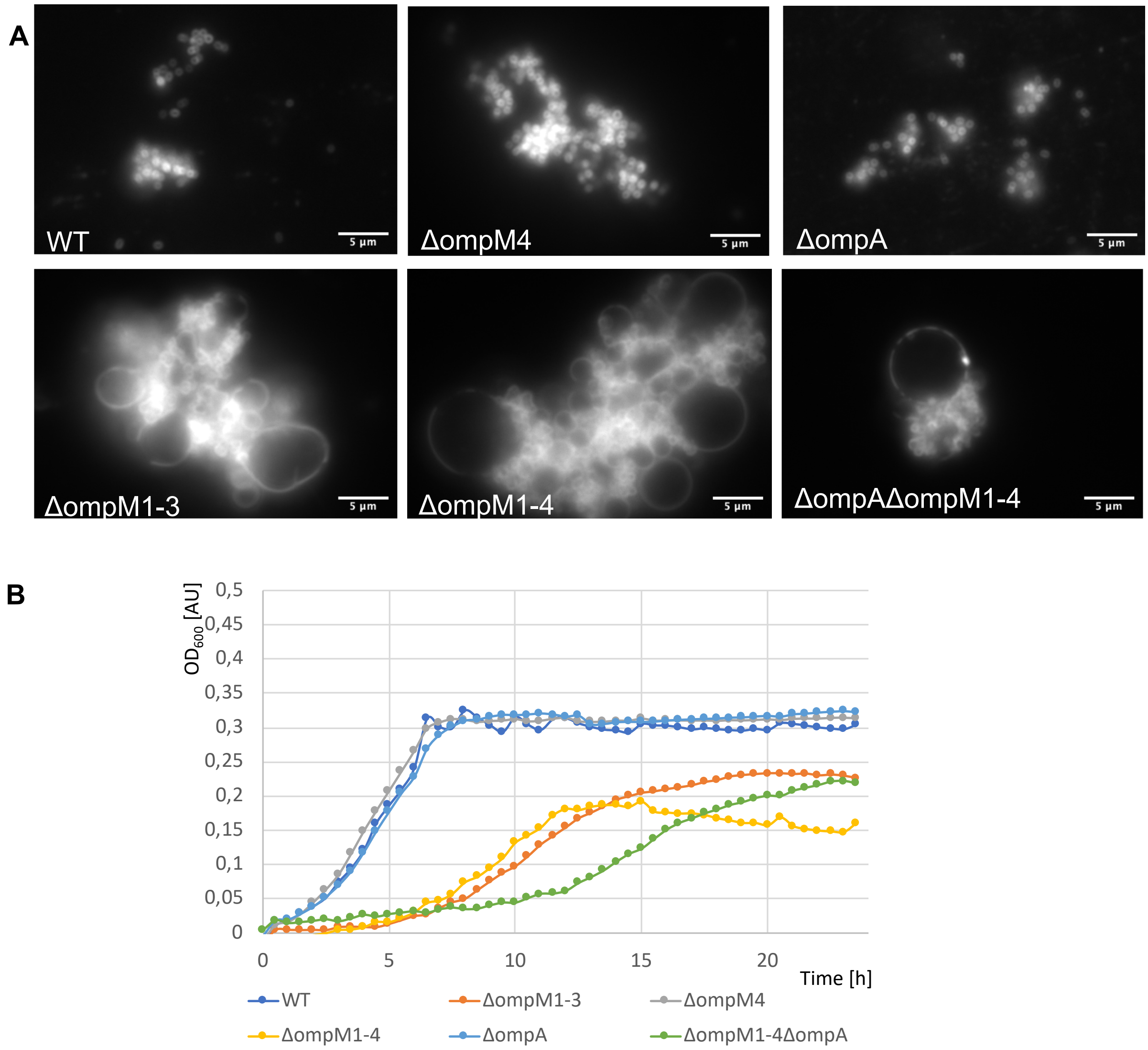


**Supplementary Figure S3. Phenotype of different *V. parvula* mutants generated in this study as compared to the WT. (A) –** Epifluorescence observation of cells labelled with the biological membrane staining dye FM 4-64 (Thermo Fisher Scientific). **(B)** – Growth curves. Cultures were made in BHILC medium in 96-well plates at 37°C under anaerobic conditions.

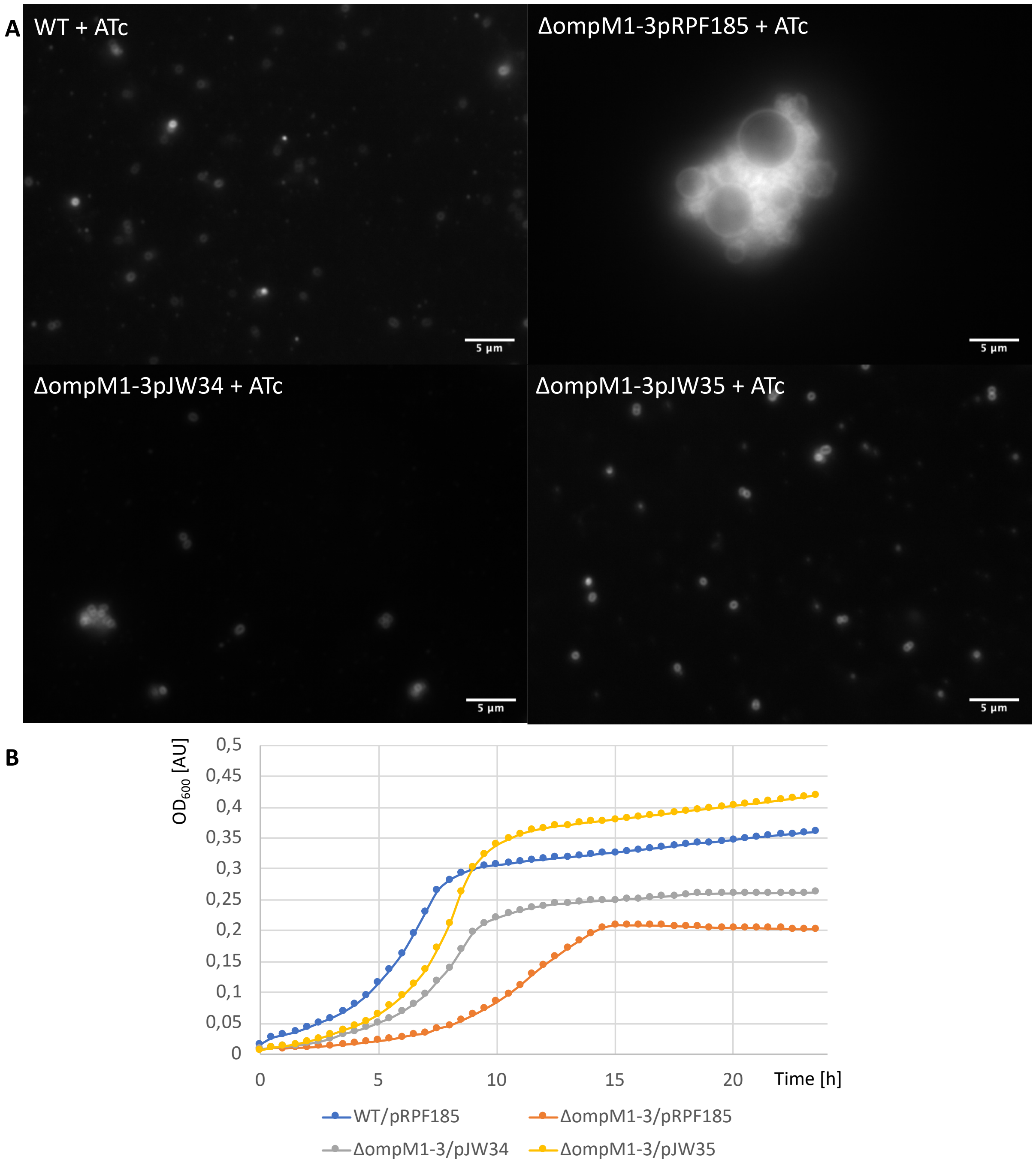


**Supplementary Figure S4. Phenotype of different *V. parvula* mutants generated in this study as compared to the WT. (A) –** Epifluorescence observation of cells labelled with the biological membrane staining dye FM 4-64 (Thermo Fisher Scientific). **(B)** – Growth curves. Cultures were made in BHILC medium in presence of 250 $\mu$g/l of anhydrotetracycline in 96-well plates at 37°C under anaerobic conditions. pRPF185 – empty vector, pJW34 – a vector expressing native OmpM1 under the control of *tet* promoter, pJW35 – a vector expressing HA-tagged OmpM1 under the control of *tet* promoter

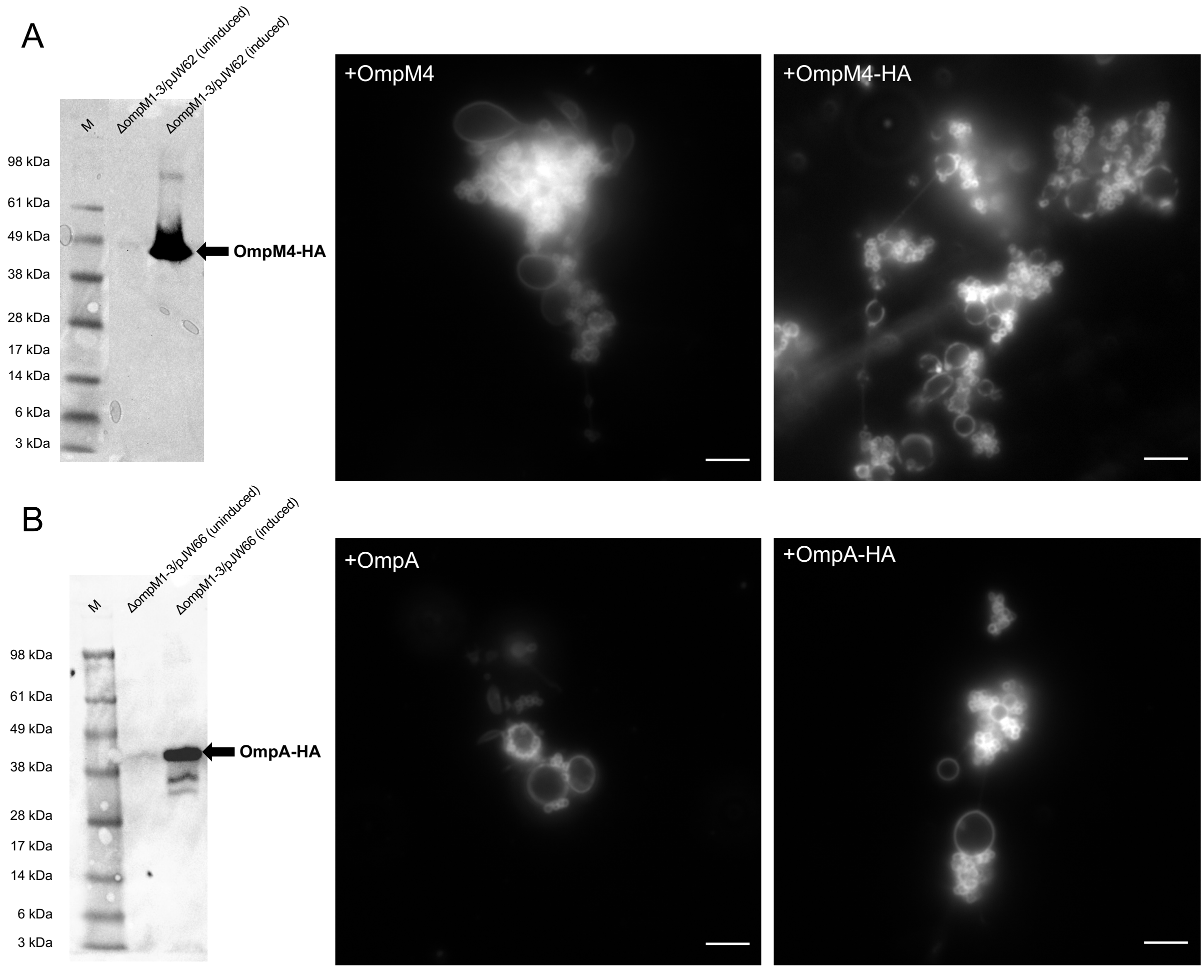


**Supplementary Figure S5.** Western blot control and FM 4-64 epifluorescence images of unsuccessful complementation of the *ompM1-3* mutant by overexpressing OmpM4 (**A**) or OmpA from plasmid (**B**). For each figure the left panel shows the anti-HA tag western blot, the middle panel the epifluorescence images of expression of the untagged version (from pJW63 for OmpM4 and pJW65 for OmpA) and the right panel the epifluorescence images of expression of the HA-tagged version (from pJW62 for OmpM4 and pJW66 for OmpA). The protein expression was induced overnight with 250 $\mu$g/l of anhydrotetracycline. Scale bars represent 5 $\mu$m.

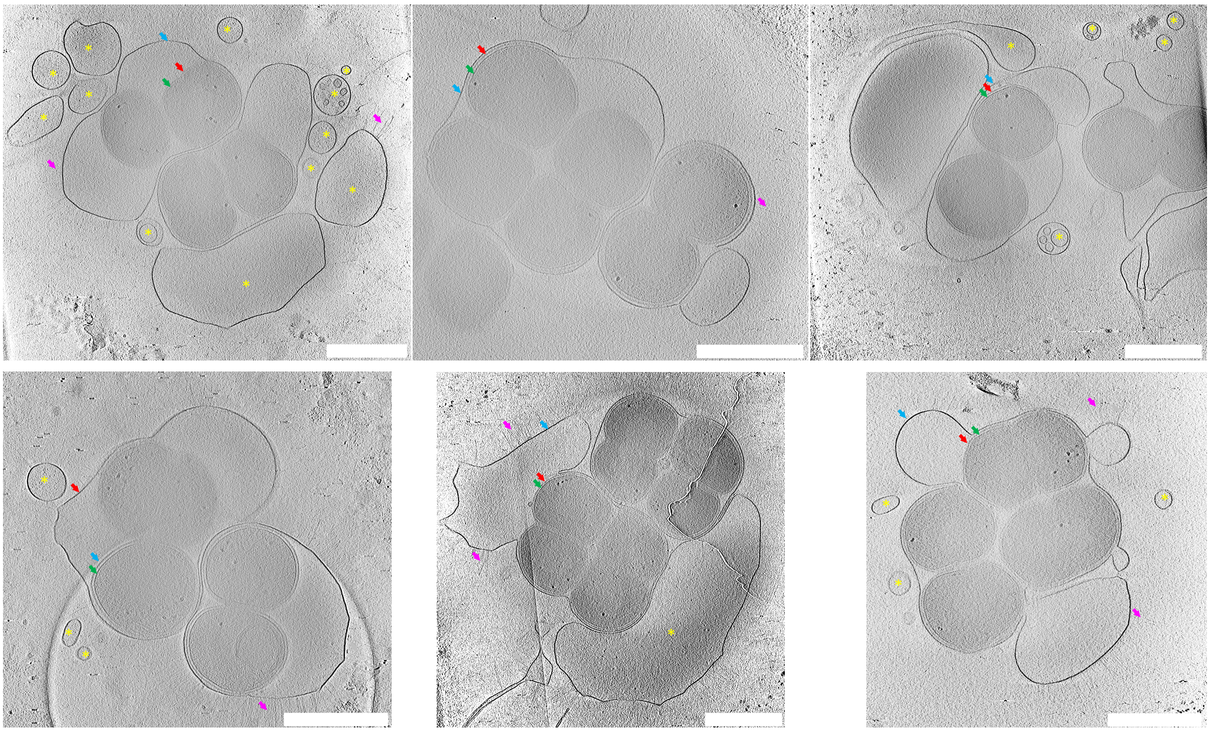

**Supplementary Figure S6** – slices of cryoelectron tomography of different positions of a mesh grid containing the *ΔompM1-3* mutant. Scale bars represent 0.5 $\mu$M, blue arrows OM, red arrows peptidoglycan, green arrows IM, grey arrows fimbriae, yellow asterisks indicate empty vesicles.

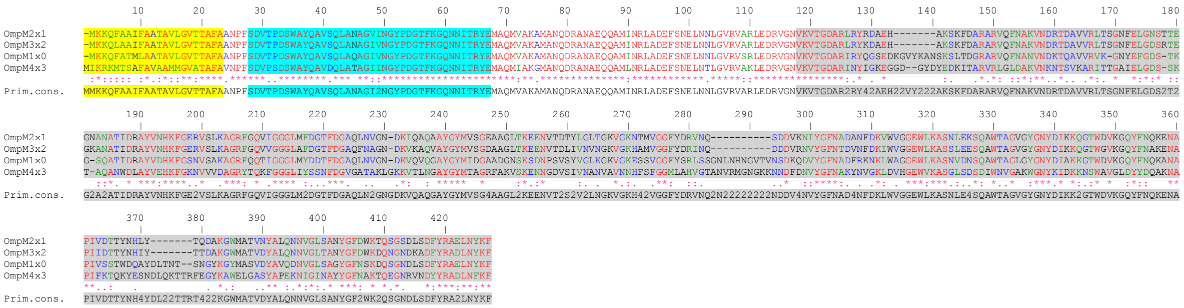

**Supplementary Figure S7. The ClustalW (Thompson et al., 1994) alignment of proteic sequences of the OmpM.** The domains were indicated by highlighting: yellow – signal peptide, cyan – S-layer homology domain, grey – beta barrel. 100% identity is indicated by asterisks and red letters, strong similarity by colons and green letters, weak similarity by points and blue letters, the primary consensus sequence is given below.

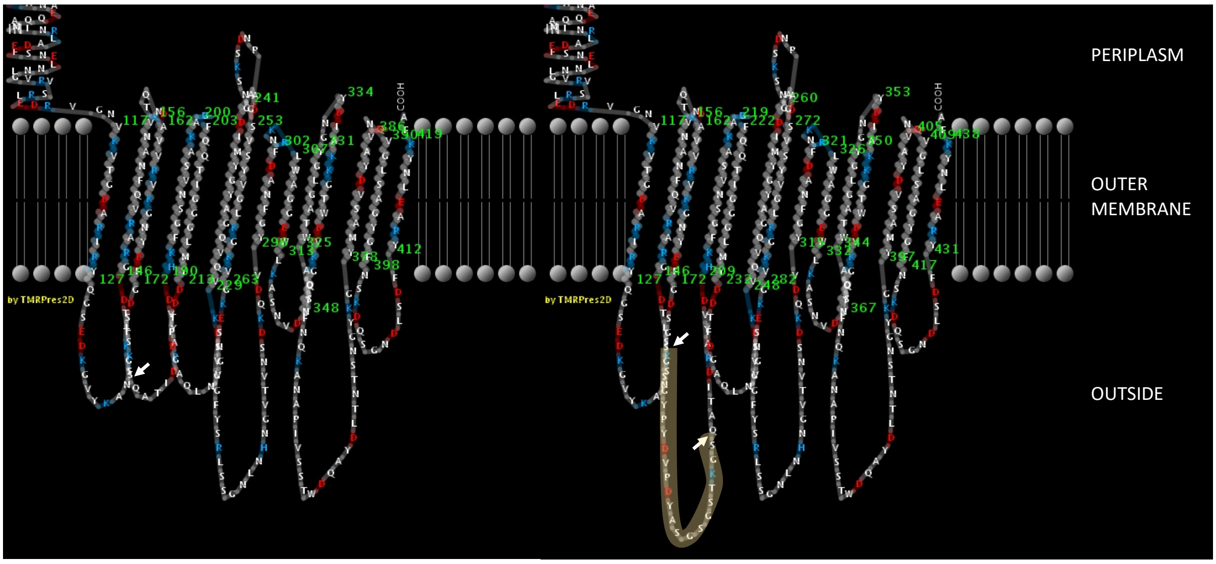


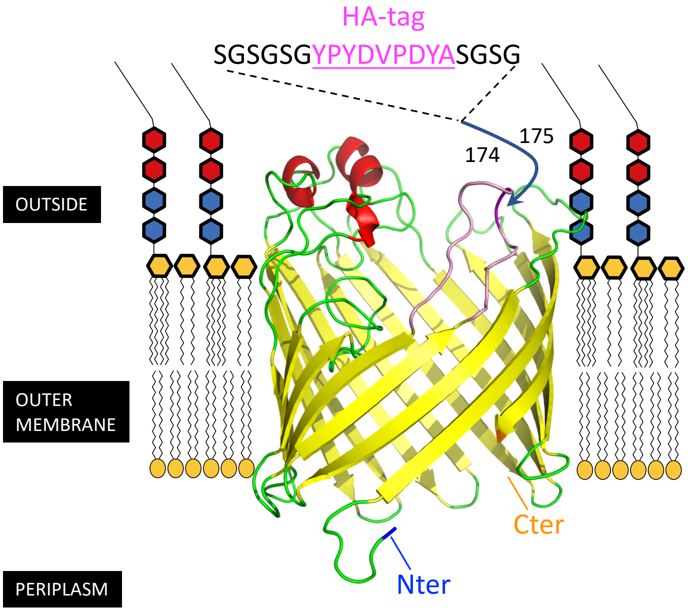


**Supplementary figure S8.** Top: TMRPres2D* prediction of the transmembrane beta strands and extracellular loops for the native (left) and HA-tagged (right) OmpM1. The position of the HA-tag insertion is marked by arrows (and the HA-tag with the linker is highlighted in pale yellow). Bottom: Phyre** prediction of the beta barrel part of OmpM protein (residues 109-420) with the tag insertion position marked.

* http://bioinformatics.biol.uoa.gr/PRED-TMBB/input.jsp
** http://www.sbg.bio.ic.ac.uk/~phyre2/html/page.cgi?id=index

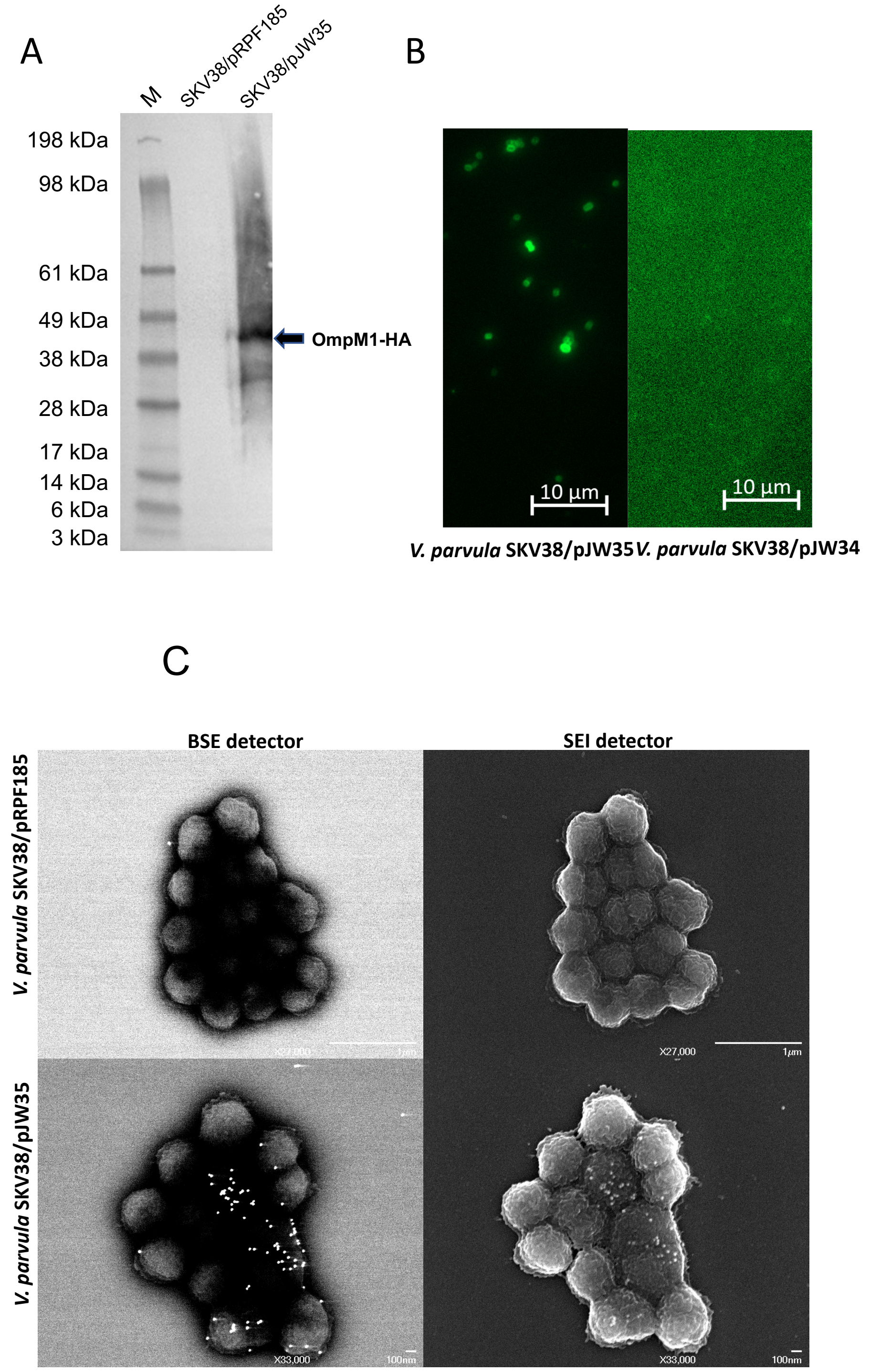


**Supplementary Figure S9**. **The expression of OmpM1 in *V. parvula*. A** – Western blot anti HA-tag. M – SeeBlue Plus Prestained Protein Standard (Invitrogen). The detected protein is marked with an arrow. **B** – Immunofluorescence images of anti-HA tag labelling on non-permeabilised cells. **C** – Scanning electron microscopy images of anti-HA immunogold (with 20 nm gold particles) indirect detection on non-permeabilised cells, obtained with a backscattered electrons detector (BSE) to detect the gold labelling and with a secondary electron detector (SEI) to image the surface of the sample. Comparison of pRPF185 (empty vector, pJW34), a vector expressing untagged OmpM1, and pJW35, a vector expressing HA-tagged OmpM1 both under the control of *tet* promoter. All cultures were induced overnight with 250 $\mu g/l$ anhydrotetracycline.

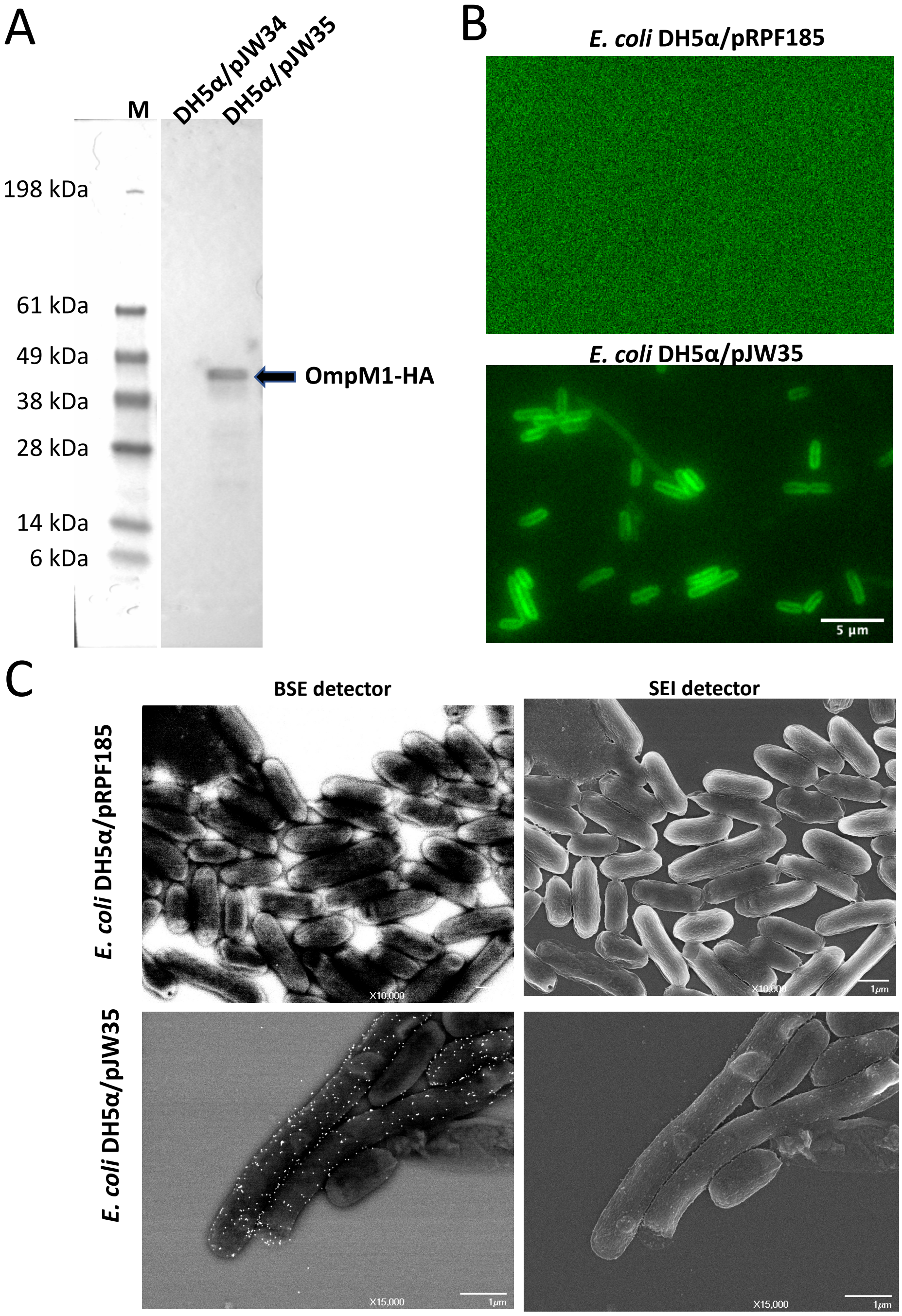


**Supplementary Figure S10**. **The expression of OmpM1 in *E. coli*. A** – Western blot anti HA-tag. M – SeeBlue Plus Prestained Protein Standard (Invitrogen). The detected protein is marked with an arrow. **B** – Immunofluorescence images of anti-HA tag labelling on non-permeabilised cells. **C** – Scanning electron microscopy images of anti-HA immunogold (with 15 nm gold particles) indirect detection on non-permeabilised cells, obtained with a backscattered electrons detector (BSE) to detect the gold labelling and with a secondary electron detector (SEI) to image the surface of the sample. Comparison of pRPF185 (empty vector, pJW34), a vector expressing untagged OmpM1, and pJW35, a vector expressing HA-tagged OmpM1 both under the control of *tet* promoter. All cultures were induced overnight with 250 $\mu$g/l of anhydrotetracycline.

### Supplementary Tables

| Strain | Genotype | Reference or Source |
|---|---|---|
| ***E. coli* DH5α** | *F- endA1 glnV44 thi-1 recA1 relA1 gyrA96 deoR nupG purB20* φ80d*lacZ*ΔM15 Δ(*lacZYA-argF*)U169, hsdR17($r_K^- m_K^+$), $\lambda^-$ | Promega |
| ***E. coli* MFDpir** | *MG1655 RP4-2-Tc ::[Δmu1 ::aac(3)IV-ΔaphA-Δnic35-Δmu2 ::zeo] ΔdapA :☹erm-pir) ΔrecA* | (Ferrières et al., 2010) |
| ***V. parvula* SKV38** | Wild type strain | (Knapp et al., 2017) |
| ***V. parvula* SKV38 3F7** | *ΔFNLLGLLA_00516::ermE* | (Béchon et al., 2020) |
| ***V. parvula* SKV38 ΔompM4** | *ΔFNLLGLLA_00231::catP* | This study |
| ***V. parvula* SKV38 ΔompM1-3** | *ΔFNLLGLLA_01389-7::tetM* | This study |
| ***V. parvula* SKV38 ΔompM1-4** | *ΔFNLLGLLA_00231::catP ΔFNLLGLLA_01389-7::tetM* | This study |
| ***V. parvula* SKV38 ΔompA** | *ΔFNLLGLLA_00518::catP* | This study |
| ***V. parvula* SKV38 ΔompAΔompM1-4** | *ΔFNLLGLLA_00231::ermE ΔFNLLGLLA_00231::catP ΔFNLLGLLA_01389-7::tetM* | This study |

**Supplementary Table S1. Bacterial strains used in this study.**

| Plasmid | Description | Reference or Source |
|---|---|---|
| **pBJSL2** | Source of tetracycline cassette | (Liu et al., 2012) |
| **pJW03** | pUC18 with the upstream fragment of *FNLLGLLA_01389* (*ompM1*), the *tetM* and the downstream fragment of *FNLLGLLA_01387* cloned together at the *Xba*I site | This study |
| **pJW34** | pRPF185 with native *FNLLGLLA_01389* with its native RBS cloned in *SacI* site | This study |
| **pJW35** | pRPF185 with HA-tagged *FNLLGLLA_01389* (*ompM1*) with its native RBS cloned in *SacI* site | This study |
| **pJW50** | pRPF185 with HA-tagged *FNLLGLLA_00231* (*ompM4*) with its native RBS cloned in *SacI* site | This study |
| **pJW62** | pRPF185 with HA-tagged *FNLLGLLA_00231* (*ompM4*) with *FNLGLLA_01389*'s RBS cloned in *SacI* site | This study |
| **pJW63** | pRPF185 with native *FNLLGLLA_00231* (*ompM4*) with *FNLGLLA_01389*'s RBS cloned in *SacI* site | This study |
| **pJW65** | pRPF185 with native *FNLLGLLA_00518 (ompA)* with *FNLGLLA_01389*'s RBS cloned in *SacI* site | This study |
| **pJW66** | pRPF185 with HA-tagged *FNLLGLLA_00518 (ompA)* with *FNLGLLA_01389*'s RBS cloned in *SacI* site | This study |
| **pRPF185** | *Escherichia/Clostridium* shuttle conjugative expression vector with *tet* promoter | (Fagan & Fairweather, 2011) |
| **pUC18** | *E. coli* cloning vector, $Amp^r$ | Thermo Fischer Scientific |

**Supplementary Table S2. Plasmids used in this study**

| Primer | Sequence | Description |
|---|---|---|
| **JW21** | AGTCGGATAGATAAAGTACG | Reverse primer for verification of the upstream junction in *ΔompM1-3* mutant |
| **JW22** | CGGTTATATCGAGCTATCAT | Forward primer for verification of the downstream junction in *ΔompM1-3* mutant |
| **JW49** | TGGATCAAGTAACAACAAAACAGC | FNLLGLLA_00231 upstream fragment – forward primer |
| **JW50** | ttccttgatttaagccccgACCTCACACACGAATAATAACAAGA | FNLLGLLA_00231 upstream fragment – reverse primer for assembly with *catP* resistance marker |
| **JW51** | gaacatgtgagcaaaaggccAGTAGAATGCTATAGGGTGAGAGTGTT | FNLLGLLA_00231 downstream fragment – forward primer for assembly with *catP* resistance marker |
| **JW52** | AGAACAGATTTTTACTCTTACCTCAAA | FNLLGLLA_00231 downstream fragment – reverse primer |
| **JW53** | CGGGGCTTAAATCAAGGAA | *catP* – forward primer |
| **JW54** | GGCCTTTTGCTCACATGTTC | *catP* – reverse primer |
| **JW56** | GCTACTCCATCGTCATTAT | Reverse primer for verification of the downstream junction in *ΔompM1-3* mutant |
| **JW57** | TCACAACTACAAGGGGCTTTTT | Forward primer for verification of the upstream junction in *ΔompM4* mutant |
| **JW58** | TTCGTTTCCAAGAAGAATCCA | Reverse primer for verification of the downstream junction in *ΔompM4* mutant |
| **JW59** | GGATTTCACATTTGCCGTTT | Reverse primer for verification of the upstream junction in *ΔompM4* and *ΔompA* mutant |
| **JW60** | GCTTCCTCGCTCACTGACTC | Forward primer for verification of the downstream junction in *ΔompM4* and *ΔompA* mutant |

| | | |
|---|---|---|
| **JW92** | GTGATGGGGGATAATCGAAA | Forward primer for verification of the upstream junction in *ΔompM1-3* mutant |
| **JW97** | aagcttgcatgcctgcaggtcgactCCGAGACGGCTAGGTTGAATC | FNLLGLLA_01389 upstream fragment – forward primer for cloning into pUC18 *Xba*I site by Gibson assembly - |
| **JW98** | cctgtggatccacTTGTGCCAATTGGGATACTG | FNLLGLLA_01389 upstream fragment – reverse primer for Gibson assembly with *tetM* resistance marker |
| **JW99** | ccaattggcacaaGTGGATCCACAGGACACAATATC | *tetM* – forward primer for Gibson assembly with FNLLGLLA_01389 upstream fragment |
| **JW100** | agactcccgaaggAGTAAAATGCAGGCGAGTG | *tetM* - reverse primer for Gibson assembly with FNLLGLLA_01387 upstream fragment |
| **JW101** | cctgcattttactCCTTCGGGAGTCTCTTTTTG | FNLLGLLA_01387 downstream fragment - forward primer for Gibson assembly with *tetM* resistance marker |
| **JW102** | ttcgagctcggtacccggggatcctTGACACATTAGGGGCAATG | FNLLGLLA_01387 downstream fragment – reverse primer for cloning into pUC18 *Xba*I site by Gibson assembly |
| **JW122** | ccgctgccgctaccAGAGTCACCAAATTCGTAATTAC | FNLLGLLA_01389's first part – reverse primer for Gibson assembly with an HA-tag |
| **JW123** | tggtgactctggtagcggcagcggttatccatatgatgttccagattatgctag cggtagcggcagcACTAAAGGTTCTCAAGCAAC | FNLLGLLA_01389's second part – forward primer adding HA-tag |
| **JW172** | agcgttaacagatctgagctTAAGAAAGAAGGAATTCATTATGAAAAAAC | FNLLGLLA_01389 with its native RBS – forward primer for cloning in *Sac*I site of pRPF185 by Gibson assembly |
| **JW173** | tctcctttactgcaggagctTTAGAATTTGTAATTTAATTCTGCACG | FNLLGLLA_01389 – reverse primer for cloning in *Sac*I site of pRPF185 by Gibson assembly |
| **JW209** | agcgttaacagatctgagctATTCGTGTGTGAGGTTTATTATG | FNLLGLLA_00231 with its native RBS – forward primer for cloning in *Sac*I site of pRPF185 by Gibson assembly |
| **JW210** | gctgccgctaccgctagcataatctggaacatcatatggataaccgctgccgct accGCTATCACCTAACTCGATAGC | FNLLGLLA_00231's first part – reverse primer for Gibson assembly with an HA-tag |
| **JW211b** | ggtagcggcagcggttatccatatgatgttccagattatgctagcggtagcggc agcTCTAAAACAGCACAAGCTAATTGGGATTTAGCTTATGTAGAGCAC | FNLLGLLA_00231's second part – forward primer adding HA-tag |
| **JW212b** | tctcctttactgcaggagctTTAGAATTTGAAGTTAAGATCCGCACGATAGTAA TCATTTACGCGATTGC | FNLLGLLA_00231 – reverse primer for cloning in *Sac*I site of pRPF185 by Gibson assembly |
| **JW252** | agcgttaacagatctgagcttaagaaagaaggaattcattATGATTAAACGTAA AATGACTAGTG | FNLLGLLA_00231 with FNLLGLLA_01389's RBS – forward primer for cloning in *Sac*I site of pRPF185 by Gibson assembly |
| **JW255** | agcgttaacagatctgagcttaagaaagaaggaattcattATGAATAAAAAATT ATTAGCTTTATTGGC | FNLLGLLA_00518 with FNLLGLLA_01389's RBS – forward primer for cloning in *Sac*I site of pRPF185 by Gibson assembly |
| **JW256** | tctcctttactgcaggagctTTAGCGATGAATATAAACGTCTAC | FNLLGLLA_00518 – reverse primer for cloning in *Sac*I site of pRPF185 by Gibson assembly |
| **JW257** | gctgccgctaccgctagcataatctggaacatcatatggataaccgctgccgct accGCCGAACATTTTATCATTCAAAC | FNLLGLLA_00518's first part – reverse primer for Gibson assembly with an HA-tag |
| **JW258** | ggtagcggcagcggttatccatatgatgttccagattatgctagcggtagcggc agcACTAGCTATAGTGATAGAATGCAAG | FNLLGLLA_00518 – reverse primer for cloning in *Sac*I site of pRPF185 by Gibson assembly |
| JW278 | tggtggtatgcttgctcatg | Internal forward primer for verification of FNLLGLLA_00231 presence |
| JW279 | gatagtaatcatttacgcgattg | Internal reverse primer for verification of FNLLGLLA_00231 presence |
| JW280 | ccttggctccaaagtagacg | Internal forward primer for verification of FNLLGLLA_00518 presence |
| JW281 | agacgatcggcagatacacc | Internal reverse primer for verification of FNLLGLLA_00518 presence |
| **TNT3-amont** | TATAATTCCCTCCAAAAACAGCCC | FNLLGLLA_00518 upstream fragment – reverse primer |
| **TNT3-aval** | GTTCCTTATCAAGCAACATATAAACAGC | FNLLGLLA_00518 downstream fragment – reverse primer |
| **TNT3-catP** | gacttattataggactaatctagataagtctagagaTTAACTATTTATCAATTC CTGCAATTCGTTTAC | *catP* – reverse primer for assembly with the downstream fragment of FNLLGLLA_0518 |
| **TNT3-ermE** | gacttattataggactaatctagataagtctagagaTTATTTCCTCCCGTTAAATAATAGATAAC | *ermE* – reverse primer for assembly with the downstream fragment of FNLLGLLA_0518 |
| **TNT3-INTermE** | ccctttacctgttccaatttcg | Reverse primer for verification of the upstream junction in *ΔompA* mutant |
| **TNT3-exterieur** | GTAATACCTACCATCACGCCTAG | Reverse primer for verification of the downstream junction in *ΔompA* mutant |
| **TNT5-amont** | CACTTTGATGGGCATTATGGCTAT | FNLLGLLA_00518 upstream fragment – forward primer |
| **TNT5-aval** | TCTCTAGACTTATCTAGATTAGTCCTATAATAAGTC | FNLLGLLA_00518 downstream fragment – forward primer |
| **TNT5-catP** | gggctgtttttggagggaattataATGGTATTTGAAAAAATTGATAAAAATAGT TGGAAC | *catP* – forward primer for assembly with the upstream fragment of FNLLGLLA_0518 |
| **TNT5-exterieur** | GACCAGATATTGCACGTACTTATGG | Forward primer for verification of the upstream junction in *ΔompA* mutant |
| **TNT5-ermE** | gggctgtttttggagggaattataATGAACAAAAATATAAAATATTCTCAAAAC | *ermE* – forward primer for assembly with the upstream fragment of FNLLGLLA_0518 |
| **TNT5-INTermE** | gggtcaatcgagaatatcgtcaac | Forward primer for verification of the downstream junction in *ΔompA* mutant |

**Supplementary Table S3. Primers used in this study. The part of the primer that hybridizes with the template DNA is indicated in majuscules.**

#### Supplementary Materials and Methods

##### DNA manipulations

PCR reactions for cloning applications were carried out using Phusion HiFi Master Mix (Thermo Fisher Scientific) according to manufacturer's protocol. PCR reactions for the control of constructions were carried out using the DreamTaq Green MasterMix (Thermo Fisher Scientific). All primers were provided by Merck or Eurofins. PCR products were purified using the NucleoSpin Gel and PCR Clean-up kit (Macherey-Nagel). Restriction enzymes were of the FastDigest family of products (Thermo Fischer Scientific). Digestion products were isolated on agarose gels and purified with the NucleoSpin Gel and PCR Clean-up kit (Macherey-Nagel). Plasmid isolation was performed with NucleoSpin Plasmid kit (Macherey-Nagel). Sequence in silico manipulation was carried out using SnapGene (GSL Biotech, www.snapgene.com). Primers were designed either with NEBuilder (New England Biolabs, nebuilder.neb.com) or Primer3Plus (Untergasser et al., 2007).

##### Generation of *ompM* and *ompA* deletion mutants

FNLLGLLA_00231 (*ompM4*) mutant was generated using the technique described previously (Knapp et al., 2017). Briefly, the upstream and downstream fragments were amplified by PCR using *V. parvula* SKV38 gDNA as template with, respectively, JW49/JW50 and JW51/JW52 primer couples. *catP* resistance gene was amplified using JW53/JW54 primer couple and pRPF185 plasmid as a template. JW50 and JW51 primers contained overhangs complementary to the *catP* fragment. Next, the linear construct containing the upstream and downstream fragments flanking the *catP* resistance marker was assembled by PCR using three previously generated fragments as a template and JW49/JW52 primer couple. The transformation using natural competence of *V. parvula* SKV38 strain and subsequent double homologous recombination was performed following the protocol described previously (Knapp et al., 2017), yielding Δ*ompM4* strain. The correct mutant construction was verified by PCR amplification (and sequencing of amplicons) of the recombination junction zones.

FNLLGLLA_00518 (*ompA*) mutant was generated using the same approach. The upstream and downstream fragments were amplified by PCR using *V. parvula* SKV38 gDNA as template with, respectively, TNT5-amont/TNT3-amont and TNT5-aval/TNT3-aval primer couples. The *catP* resistance gene was amplified using TNT5-catP/TNT3-catP primer couple and pRPF185 plasmid as a template. TNT5-catP and TNT3-catP primers contained overhangs complementary to the, respectively, upstream and downstream fragments. Next, the linear construct containing the upstream and downstream fragments flanking the *catP* resistance marker was assembled by a PCR using three previously generated fragments as a template and TNT5-amont/TNT3-aval primer couple. The transformation using natural competence of *V. parvula* SKV38 strain and subsequent double homologous recombination was performed following the protocol described previously (Knapp et al., 2017), yielding the Δ*ompA* strain. The correct mutant construction was verified by PCR amplification (and sequencing of amplicons) of the recombination junction zones.

For FNLLGLLA_01387-9 (*ompM1-3*) deletion, the PCR assembly technique was inefficient. The upstream, downstream and *tetM* fragments generated by PCR using, respectively, JW97/JW98, JW101/JW102 and JW99/JW100 primer couples, and *V. parvula* SKV38 gDNA (upstream/downstream) or pBSJL2 (*tetM*) as templates wouldn't assemble together by PCR using the JW97/JW102 primer couple. Instead, they were cloned together into pUC18 *Xba*I site, yielding pJW03 vector. The integrity of its insert was verified with sequencing. The vector was then used as a template for PCR with JW97/JW102 primer couple, and the obtained product was used for the natural transformation (and subsequent double crossing over) following the usual protocol (Knapp et al., 2017), yielding Δ*ompM1-3* strain. The correct mutant construction was verified by PCR amplification (and sequencing of amplicons) of the recombination junction zones.

Generation of quintuple mutant

The generation of a quintuple Δ*ompA*Δ*ompM1-4* mutant was carried out as follows. First, the upstream and downstream fragments of FNLLGLLA_0518 were amplified by PCR using *V. parvula* SKV38 gDNA as template with, respectively, TNT5-amont/TNT3-amont and TNT5-aval/TNT3-aval primer couples. The *ermE* resistance gene was amplified using TNT5-ermE/TNT3-ermE primer couple and the gDNA of *V. parvula* 3F7 strain as template. TNT5-ermE and TNT3-ermE primers contained overhangs complementary to the, respectively, upstream and downstream fragments. Next, the linear construct containing the upstream and downstream fragments flanking the *ermE* resistance marker was assembled by PCR using three previously generated fragments as a template and TNT5-amont/TNT3-aval primer couple. The transformation using natural competence of *V. parvula* SKV38 strain and subsequent double homologous recombination was performed following the protocol described previously (Knapp et al., 2017) on the Δ*ompM4* mutant, yielding Δ*ompA*Δ*ompM4* strain, which was in turn transformed by the gDNA of the Δ*ompM1-3* strain, yielding the desired Δ*ompA*Δ*ompM1-4* strain. The correct mutant construction was verified by PCR amplification (and sequencing of amplicons) of the recombination junction zones as well as the absence of *ompA* and *ompM4* genes by PCR.

OmpM1, OmpM4 and OmpA Expression vector construction

For the vector expressing the native OmpM1, the FNLLGLLA_01389 with the native RBS was amplified by PCR with JW172/JW173 primers. The obtained PCR product was cloned into *Sac*I site of pRPF185, yielding pJW34 vector. The integrity of the construct was verified by sequencing.

For the vector expressing HA-tagged OmpM1, the first part of FNLLGLLA_01387 with the native RBS was amplified by PCR with JW172/JW122 primers. The second part was amplified with JW123/JW173 primers (JW123 adding the HA-tag). *V. parvula* SKV38 gDNA was used as a template for both reactions. Next the two fragments were cloned together into the pRPF185 vector linearized by *Sac*I restriction digestion, yielding pJW35 vector. The integrity of the construct was verified by sequencing. The correct expression of HA-tagged OmpM1 protein was verified by Western Blot and immunomarking (Supplementary Figure S9). Additionally, the correct expression and addressing to the cell surface was also verified in *E. coli* (Supplementary Figure S10).

For the vector expressing HA-tagged OmpM4, the first part of FNLLGLLA_00231 with the native RBS was amplified by PCR with JW209/JW210 primers. The second part was amplified with JW211b/JW212b primers (JW211b adding the HA-tag). *V. parvula* SKV38 gDNA was used as a template for both reactions. Next, the two fragments were cloned together into the pRPF185 vector linearized by *Sac*I restriction digestion, yielding pJW50 vector. The integrity of the construct was verified by sequencing. The expression of the HA-tagged OmpM4 protein was verified by Western Blot. As the expression levels were extremely low, the HA-containing construct was amplified by PCR from pJW50 vector with JW252/JW212b primer couple, JW252 replacing the native RBS with FNLLGLLA_01389's RBS. The amplicon was then cloned into the *Sac*I site of pRPF185 vector, yielding pJW62 plasmid. The integrity of the construct was verified by sequencing. The correct expression of HA-tagged OmpM4 protein was verified by Western Blot (Supplementary Figure S5A).

For the vector expressing the native OmpM4, the FNLLGLLA_00231 was amplified by PCR with JW252/JW212b primers (JW252 adding the

FNLLGLLA_01389's RBS). The obtained PCR product was cloned into *Sac*I site of pRPF185, yielding pJW63 vector. The integrity of the construct was verified by sequencing.

For the vector expressing the native OmpA, the FNLLGLLA_00518 was amplified by PCR with JW255/JW256 primers (JW255 adding the FNLLGLLA_01389's RBS). The obtained PCR product was cloned into *Sac*I site of pRPF185, yielding pJW65 vector. The integrity of the construct was verified by sequencing.

For the vector expressing HA-tagged OmpA, the first part of FNLLGLLA_00518 was amplified by PCR with JW255/JW257 primers (JW255 adding the FNLLGLLA_01389's RBS). The second part was amplified with JW258/JW256 primers. The HA-tag was added by the complementary parts of JW257 and JW258 primers. *V. parvula* SKV38 gDNA was used as a template for both reactions. Next, the two fragments were cloned together into the pRPF185 vector linearized by *Sac*I restriction digestion, yielding pJW66 vector. The integrity of the construct was verified by sequencing. The correct expression of HA-tagged OmpA protein was verified by Western Blot (Supplementary Figure S5B).

<u>Western blot</u>

Bacterial cells were lysed by resuspension in 1x Laemmli buffer (Thermo Fisher Scientific) and heating for 10 min at 95°C, sonicated on ice, and then heated again. The gel electrophoresis was performed on BOLT Bis-Tris Plus 4-12% gel (Invitrogen) in NuPAGE MES SDS running buffer (Invitrogen) at 100 V. The proteins were transferred to a nitrocellulose membrane in TruPAGE Transfer Buffer (Invitrogen) at 10 V during 1 h. The membrane was briefly washed in water, then incubated in 5% skimmed milk (Roth or Régilait) dissolved in phosphate buffered saline with 0.5 % Tween-20 (PBST) for 1h. Two 5 min washes in PBST were made, then the membrane was incubated in a rabbit anti-HA-tag polyclonal gamma immunoglobulin (Novus Biologicals) diluted at

800 μg/l in PBST containing 1 % milk for 1h. Four 5 min washes were made in PBST, as well as a brief wash in water, and the western blot was revealed by ECL Amersham (GE Healthcare) kit.

Immunolabelling, epifluorescence and scanning electron microscopy (SEM)

A protocol for marking the outer membrane proteins without permeabilization was used. Bacterial liquid culture was washed twice with phosphate buffered saline (PBS) and resuspended in an equal volume of PBS. The bacterial suspension was deposited onto a poly-L-lysine coated microscopy slide, incubated 5 min, fixed with 4% paraformaldehyde in PBS for 10 min (for SEM observation 0.05% of glutaraldehyde was added to the fixing solution), quenched for 3 min with 50 mM ammonium chloride solution in PBS, washed three times with PBS, then the non-specific sites were saturated by 15 min incubation in 0.5% bovine serum albumin (BSA) solution in PBS.

Next, for immunofluorescence the cells were incubated for 45 min with 100 μg/l of rat monoclonal anti-HA-tag gamma immunoglobulin coupled to fluorescein (Roche) diluted in PBS containing 0.5% BSA. After three PBS washes and one water wash, cells were mounted with Dako fluorescent mounting medium and observed using an AxioPlan 2 (Carl Zeiss) microscope equipped with AttoArc HBO 100W light source (Carl Zeiss).

For SEM, the cells were incubated for 45 min with a mouse anti-HA-tag monoclonal gamma immunoglobulin (Thermo Fisher Scientific) diluted at 2 mg/l in PBS containing 0.5 % BSA. Three washes with PBS were performed and the cells were incubated for another 45 min with a goat anti-mouse gamma immunoglobulin conjugated to 20 nm gold beads (BBInternational) diluted at 4 mg/l in PBS containing 0.5 % BSA. After three PBS washes and one water wash, the gold immunolabelled samples were air-dried, then post-fixed for 30 min in 2.5% glutaraldehyde in 0.2 M sodium cacodylate buffer. Three washes were performed in 0.2 M sodium cacodylate.

Cells were then progressively dehydrated by 5 min serial immersions in 25%, 50%, 75% and 95% ethanol and by two 10 min incubation in anhydrous ethanol, followed by critical point drying with a Leica EM CPD 300. Finally, dried specimens were sputter-coated with a 10 nm conductive carbon layer using a gun ionic evaporator PEC 682. The samples were imaged in a JEOL JSM 6700F field emission scanning electron microscope operating at 7kV using a secondary electron detector (SEI) and a backscattered electron detector (YAG) to detect gold immunolabelling.
